# A General LSTM-based Deep Learning Method for Estimating Neuronal Models and Inferring Neural Circuitry

**DOI:** 10.1101/2021.03.14.434027

**Authors:** Kaiwen Sheng, Peng Qu, Le Yang, Xiaofei Liu, Liuyuan He, Youhui Zhang, Lei Ma, Kai Du

**Affiliations:** Institute for Artificial Intelligence, Peking University, Beijing, China; Beijing Academy of Artificial Intelligence, Beijing, China; School of Electronics Engineering and Computer Science, Peking University, Beijing, China; Department of Computer Science and Technology, Tsinghua University, Beijing, China

## Abstract

Computational neural models are essential tools for neuroscientists to study the functional roles of single neurons or neural circuits. With the recent advances in experimental techniques, there is a growing demand to build up neural models at single neuron or large-scale circuit levels. A long-standing challenge to build up such models lies in tuning the free parameters of the models to closely reproduce experimental recordings. There are many advanced machine-learning-based methods developed recently for parameter tuning, but many of them are task-specific or requires onerous manual interference. There lacks a general and fully-automated method since now. Here, we present a Long Short-Term Memory (LSTM)-based deep learning method, General Neural Estimator (GNE), to fully automate the parameter tuning procedure, which can be directly applied to both single neuronal models and large-scale neural circuits. We made comprehensive comparisons with many advanced methods, and GNE showed outstanding performance on both synthesized data and experimental data. Finally, we proposed a roadmap centered on GNE to help guide neuroscientists to computationally reconstruct single neurons and neural circuits, which might inspire future brain reconstruction techniques and corresponding experimental design. The code of our work will be publicly available upon acceptance of this paper.

## Introduction

As the recent progress in experimental techniques, such as patch-seq(Cadwell et al., 2016), calcium imaging(Grienberger & Konnerth, 2012), and the combination of two-photon calcium imaging and electron microscopy(Briggman et al., 2011), neuroscientists are able to observe the connectivity and activities of neurons in a larger scale. Such progress brings forward a new challenge for the neuroscience community, namely not only to uncover the computational principles limited at single neuron or microcircuits levels, but also at a large-scale circuit level. Since now, computational models constrained by experimental findings are one of the most popular tools to describe the underlying computational mechanisms(Hodgkin & Huxley, 1952; Mainen & Sejnowski, 1996; Koch & Segev, 1998; Abbott, 1999; Izhikevich, 2003; Pillow et al., 2008; Adesnik et al., 2012; Markram et al., 2015; Du et al., 2017; Teeter et al., 2018; Atiya et al., 2019; Billeh et al., 2020; Egger et al., 2020; Hjorth et al., 2020; Oesterle et al., 2020). However, there are a large number of free parameters in the models which are not constrained by experiments, and therefore it gradually becomes infeasible for neuroscientists to hand-tune those parameters as the scale of the models grows.

Until now, there have been many methods developed to automate the parameter tuning procedure (See Discussion Related Work for details). However, some methods are specifically designed for single neuron inference(Bush et al., 2005; Q. J. Huys et al., 2006; Q. J. Huys & Paninski, 2009; Pozzorini et al., 2015; Ladenbauer et al., 2019) or neural connectivity estimation(Kobayashi et al., 2019; Ladenbauer et al., 2019; Endo et al., 2020) alone. Sequential neural posterior estimation (SNPE)-based algorithms(Gonçalves et al., 2020) and some search-based algorithms are task-independent, while they require massive and careful feature engineering on raw neural responses, which is time-consuming and non-trivial. These feature-based tuning algorithms might not be suitable to computationally reconstruct large-scale neural systems. It was only recently that deep learning, which has shown tremendous ability in automatic feature extraction, has been formally proposed in single neuron inference(Ben-Shalom et al., 2019). It uses the convolutional neural network (CNN) as the feature extractor, while it treats the neural responses at each time step as independent events ignoring the dependencies along the temporal dimension. Also, its ability to infer neural circuitry and its performance on experimental data remains unclear.

To date, there lacks an efficient and general method to unify from inferring single neuronal properties, such as the conductance of ion channels, to the connectivity of large-scale neural circuits. Here, we propose the general neural estimator (GNE), which includes an LSTM component to extract temporal features of neural responses automatically. LSTM(Hochreiter & Schmidhuber, 1997) is one of the most prevalent variations of recurrent neural networks (RNN). It has been widely used in natural language processing(Sutskever et al., 2014; Brown et al., 2020), time series analysis(Gamboa, 2017). It alleviates the limitations of traditional RNN variations, namely the gradient vanishing problem, by introducing gating mechanisms, which enable it to learn long-term dependencies. Benefited from LSTM, GNE outperformed many advanced methods recently proposed, including MOO method(Druckmann et al., 2007), CNN-based method(Ben-Shalom et al., 2019), SNPE-based method(Gonçalves et al., 2020), I&F method(Ladenbauer et al., 2019), and GLMCC method(Kobayashi et al., 2019), covering a wide range of models including phenomenal models and biophysically detailed models. GNE can infer single neurons and neural circuits from raw experimental recordings (membrane potential or spike trains), which is easy to apply and eliminates onerous feature engineering. In this paper, we demonstrate that GNE replicates experimental dynamics of over 20 different mousy spiny cells from Allen Institute Cell Types Database(Allen Cell Types Database, 2016) in a single pass, and we also equip GNE with a statistical calibration component to fine-tune on some failed cases. Furthermore, GNE can precisely estimate the connectivity strengths of a neural circuit composed of 100 neurons from highly variational and noisy neural responses. Theoretically, GNE is applicable to neural circuits or neural networks of arbitrary sizes.

Based on GNE, we also present a roadmap to facilitate neuroscientists on computational reconstruction, from single neurons to neural circuitry. Unlike previous brain reconstruction, especially synaptic reconstruction, based mainly on statistics(Markram et al., 2015; Billeh et al., 2020), we showed in this paper that GNE can precisely infer the conductance of NMDA-Receptor (NMDAR) and AMPA-Receptor (AMPAR) from raw electrophysiological data, within a neural microcircuit built up by previously well-reconstructed single neuron models. Therefore, GNE provides a novel and unified procedure for reconstructing detailed biophysical neuronal models and neural circuits on a finer and more precise scale, which might shed light on the future experimental design of brain reconstruction.

## Results

### GNE Workflow and Reconstruction Roadmap

In this paper, we propose the GNE and a roadmap based on it to computationally reconstruct single neurons and neural circuits (Figure 1a). Overall, single neurons and neural circuits are computationally reconstructed by constraining model parameters by the GNE. In particular, neural circuit model can be built upon phenomenal models, such as integrate-and-fire (I&F) models or previously well-reconstructed biophysical neuronal models. The connectivity properties of the circuit models are estimated by the GNE then. The method workflow to estimator parameters of computational models is unified for both single neuronal models and neural circuit models. The workflow is composed of 4 components (Figure 1b): model hypothesis, synthesized dataset, GNE and statistical calibration. First, we need to propose a computational model for single neurons or neural circuits. The models need to be able to replicate experimental recordings. Then, we use the models to synthesize a dataset for training and evaluation. We need to manually define a prior parameter distribution and register stimulus protocols. By applying the sampled parameters from the prior distribution and stimulus protocols to the computational models, we can obtain the corresponding model responses. The sampled parameters and the model responses together form a piece of data. The synthesized dataset is then used for training the GNE. During the training procedure, an error between the predicted parameters and the groundtruth will be backpropagated(Rumelhart et al., 1985) to update the weights of the deep neural network until the training loss converges and the weights of the deep neural network will be saved as a pre-trained model (for training details, see Methods). During inference, the experimental recording is first processed to extract local temporal features through convolution units. Then, the local temporal features are further processed by LSTM cells to extract temporal dependency features. Regression is finally operated on those temporal dependency features to get the parameter predictions (Figure 1c). If the prediction cannot fully replicate the experimental recordings, the statistical calibration procedure can be applied to fine-tune on specific experimental recordings to get more accurate results (Figure 1b). Therefore, a unified fully-automated method to infer single neuronal properties and neural circuit connectivity is built up. GNE requires little manual interference and no feature engineering.

**Figure 1:**
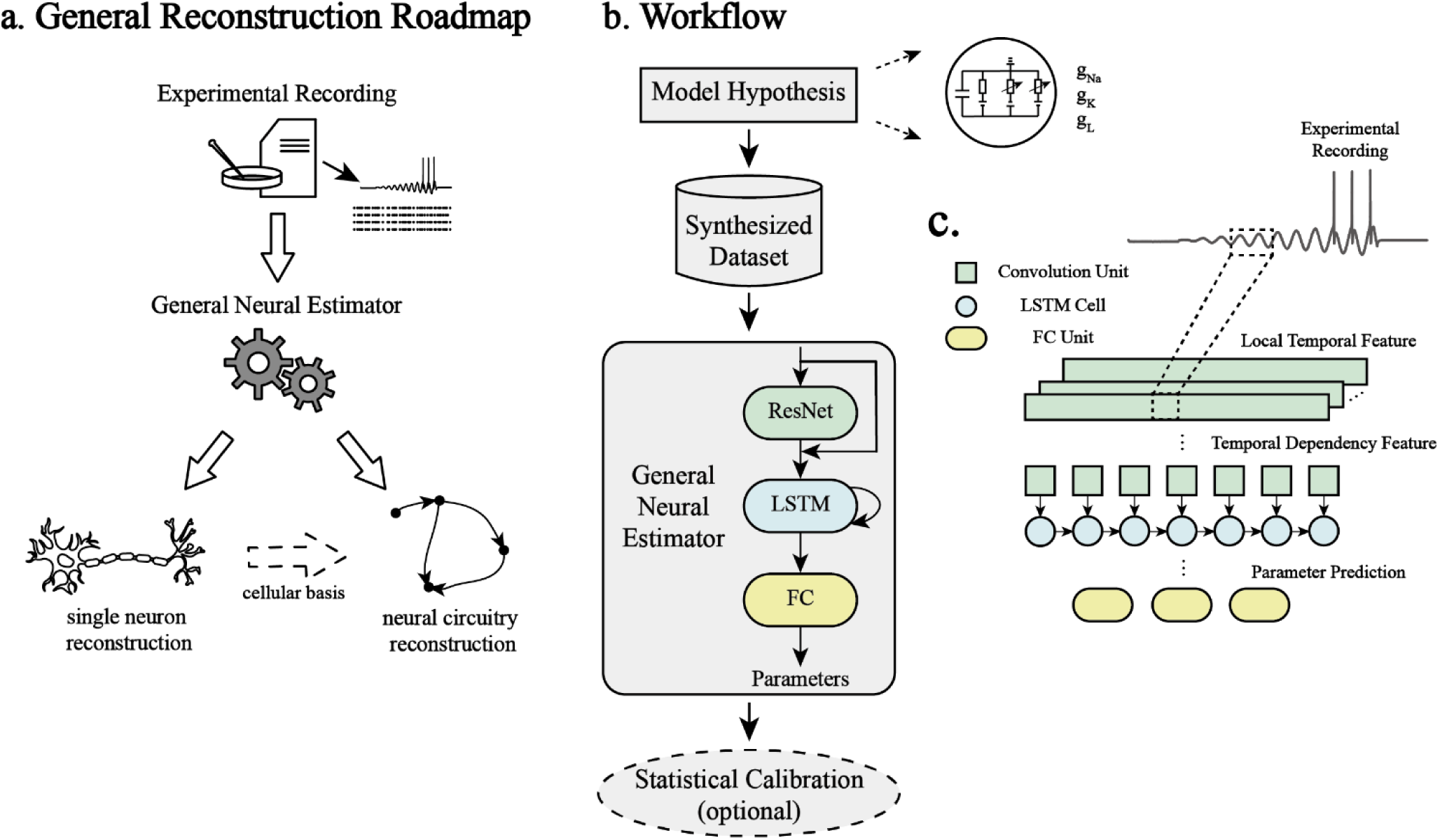
**a** Roadmap to precisely reconstruct single neurons and neural circuits. After we obtain some experimental recordings, such as intracellular and extracellular recordings, they are applied to the general neural estimator. The output of the general neural estimator can be the parameters of the computational neuronal model or the synaptic strengths within the neural circuits. To achieve neural circuitry reconstruction, there are two paths in the roadmap. First, it’s to reconstruct mathematical model of single neuronal models and then to build up biophysical neural circuitry models based on those neuronal models. Second, it’s to direct reconstruct neural circuitry from experimental recordings. **b** Workflow: first, we should manually define a computational model for single neurons and neural circuits, which we call model hypothesis. There are some free parameters in the model to be constrained by experimental recordings. For example, the three maximal conductance variables in a HH models. Then synthesized dataset is generated by applying stimulus protocols to the model and sampling parameters of the model. The dataset is used to train and evaluate the general neural estimator. After the training loss converges, experimental recordings are applied to get the predictions of the model parameters. If the performance is not accurate enough, the prediction can be further calibrated with statistical algorithms, such as SNPE, on specific experimental recordings. **c** Illustration of the deep neural network architecture. The neural responses are first processed by the convolution units to extract local temporal features and then further analyzed by LSTM to get temporal dependency features. The features are finally regressed by fully connected units to get parameter predictions.

### Deep Neural Network Architecture

#### Overall Architecture

We present a novel deep neural network architecture to perform the task of parameter estimation. The overall architecture can be divided into the following three components (Figure S1; for more details, see Methods):

1. Feature Extractor: we use a variant of ResNet-50 as the feature extractor as it has shown great performance in many fields of artificial intelligence(He et al., 2016).
2. Time Series Analyzer: we choose an LSTM(Hochreiter & Schmidhuber, 1997) as the time series analyzer because it is well known to effectively capture the temporal dynamics and dependencies.
3. Regressor: we implement 5-layer fully-connected units as the final regressor to map the feature space of raw neural responses to the parameter space.

The architecture of our design focuses on how to extract the characteristic patterns and temporal dependencies within the neural responses. Therefore, we chose ResNet-50 and LSTM as the core component of our architecture.

#### Residual Network (ResNet)

ResNet is one of the most powerful convolutional architectures to extract features nowadays. Residual Blocks (ResBlocks) are the basic components in ResNet-50 and other variants of ResNet (Figure S1b). Each ResBlock is composed of three units, where each unit contains a convolutional layer (whose kernel size is 1, 3, 1 correspondingly), a batch normalization layer, and a leaky ReLU activation layer. A skip connection associates the input with the output of the ResBlock through a summation operation. The skip connection in the ResBlock allows the network to learn the residuals, which makes it easier to train deeper neural networks easier maintaining high performance(He et al., 2016).

#### Long Short-Term Memory (LSTM)

LSTM is the most critical part of our architecture because it enables the deep neural network to capture temporal dynamics and correlations within the neural responses, rather than treating the sequence as an array of independent events, as traditional convolution does. LSTM maintains an internal loop through connections of the LSTM cells (Figure S1c). There are three gates in each LSTM cell: the forget gate, the input gate, and the output gate. The forget gate *f*_*t*_ controls whether the past cell state *C*_*t*−1_ should be forgotten. The input gate *i*_*t*_ controls what new values should be added to the cell state and the output gate *o*_*t*_ controls which part of the cell state to output. The update rules of each gate variable, cell state and hidden state are as equations (1) - (6).

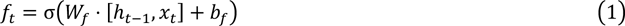

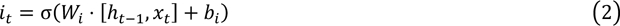

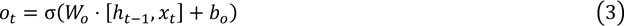

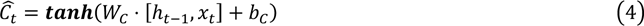

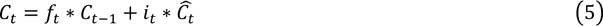

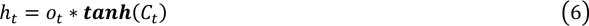

where *f*_*t*_, *i*_*t*_, *o*_*t*_ are the forget gating variable, input gating variable and output gating variable correspondingly, *W*_*f*_, *W*_*i*_, *W*_*o*_ are the forget weight, input weight and output weight correspondingly, *b*_*f*_, *b*_*i*_, *b*_*o*_ are the forget bias, input bias and output bias correspondingly, *x*_*t*_, ℎ_*t*_, *C*_*t*_ are the input, hidden state and cell state at time step *t*, and ℎ_*t*−1_, *C*_*t*−1_ are the hidden and cell state and time step *t* − 1.

### Inferring Single Neuronal Models on Synthesized Data

We trained the GNE to optimize the parameters, and compared the performance with MOO and CNN(Ben-Shalom et al., 2019) in four types of neuronal models, and we use a Hodgkin-Huxley (HH) single-compartment neuronal model as an example to illustrate our work and discoveries (see Methods for details). We first use synthesized data to train and evaluate GNE (Figure 2a), where the synthesized dataset is generated by applying stimulus on a HH single-compartment neuron model. We also demonstrate the performance comparison and noise robustness test on the HH single-compartment neuron model. Results for Izhikevich model, HH ball-stick model, L4 stellate neuron model and L5 pyramidal neuron model can be noted in Figure S2, Figure S4, Figure S6, Figure S8, Figure S10 respectively (See Supplementary for details).

**Figure 2:**
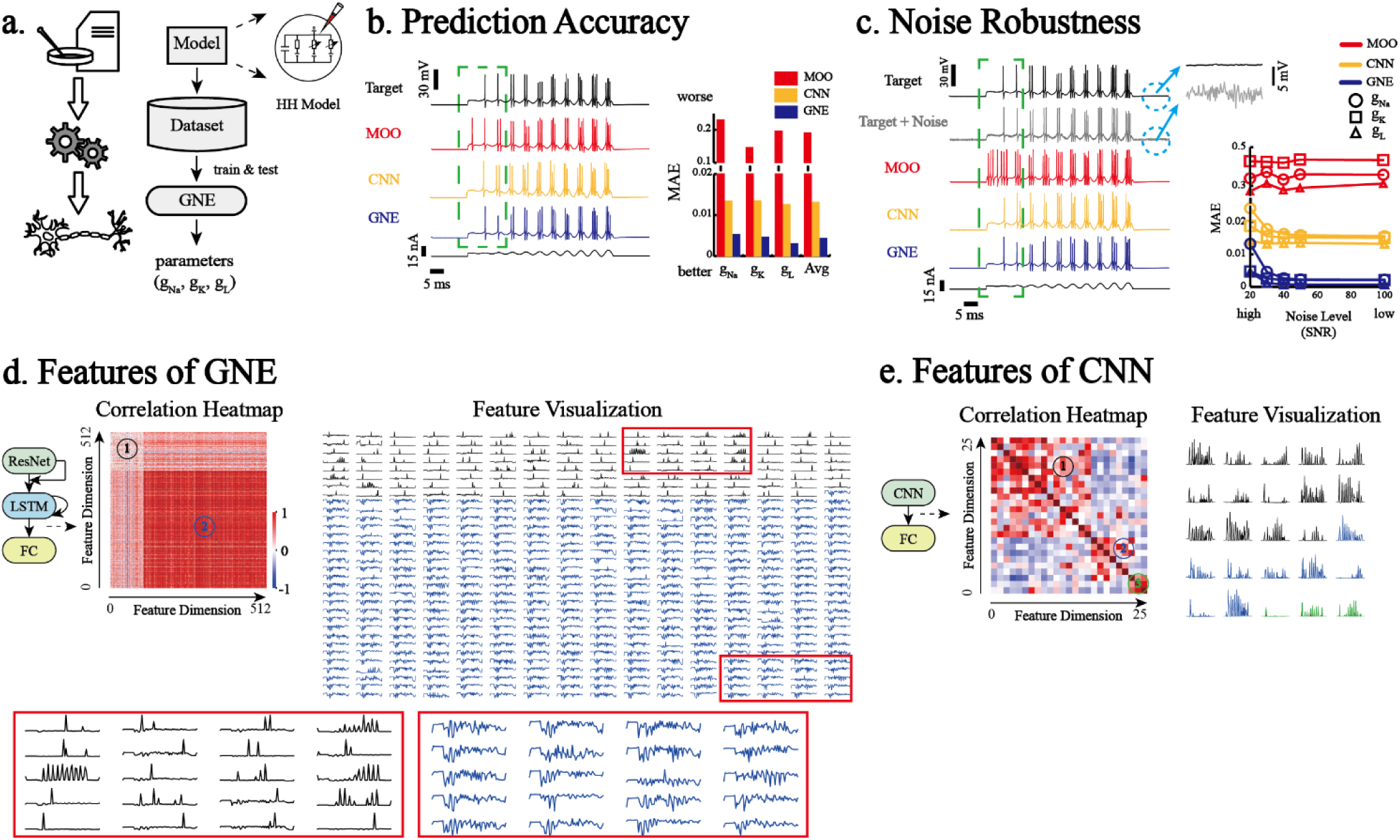
**a** Roadmap and method workflow for training and evaluation on synthesized data. First registered stimulus protocols are applied to HH single neuronal model to generate dataset for training and evaluation. The responses of the neuronal model serve as the input of GNE to get the prediction of model parameters. **b & c** Performance against MOO and CNN. Left: Examples showing response prediction under chirp stimulus protocol and step stimulus protocol, where the black curve is the groundtruth. Right: statistics of performance metric. **b** Accuracy of the predicted parameters. **c** Gaussian noise is added to the model responses at different SNR levels (20, 30, 40, 50 and 100) during searching or testing phase. The gray noisy response is added Gaussian noise with SNR level of 20. The zoom-in shows the amplificatory part of the origin model responses and noisy responses within the dashed blue circle. **d & e** Features Extracted from Deep Neural Networks before Fully Connected Layers. Left: indicates the locations of the features in the deep neural network. Middle: the correlation heatmap among features extracted from the deep neural network. Right: visualizations of the features. The color of each feature indicates the cluster it belongs to, which shares the same color as the label in the heatmap.

The dynamics of HH single-compartment neuron model are formulated as equations (7) - (8),

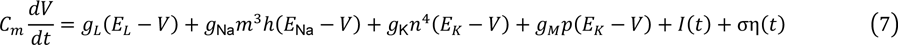

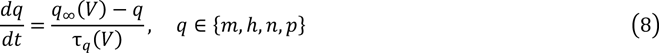

where we choose the membrane capacity *C*_*m*_ = 1μF/cm, the reversal potential *E*_Na_ = 53 mV, *E*_K_ = −107 mV and *E*_L_ = −70 mV, the initial membrane potential *d*_0_ = −70 mV, the spike threshold *d*_*T*_ = −60 mV, the density of M channel *g*_*M*_ = 0.07 mS/cm^2^, τ_max_ = 600 ms to scale the time constant through τ_*q*_(*d*), the noise factor σ = 0.1, the intrinsic noise η(*t*) as the standard Gaussian noise.

The parameters that we need to estimate are *g*_Na_, *g*_K_ and *g*_L_, whose lower bounds are [0.5, 10^−4^, 10^−4^] and upper bounds are [80, 15, 0.6]. The HH dynamics, the ranges of free parameters, and the values of other parameters are obtained from (Meliza et al., 2014; Gonçalves et al., 2020). We choose uniform distribution as the prior distribution of the parameters and chirp stimulus as the training stimulus protocol (see Methods for details of stimulus protocols).

We apply three different performance metrics to compare GNE with MOO and CNN. First, we compared the MAE between the predicted parameters and the groundtruth. GNE is 61.4, 44.3 and 93.2 times more accurate than MOO, and 3.9, 4.7 and 6.7 times than CNN in terms of predicting *g*_Na_, *g*_K_ and *g*_L_ (Figure 2b). Second, we demonstrate that the predicted parameters of GNE can well replicate the response of the target model. We show an example of model responses from the groundtruth, MOO, CNN, and GNE, on the left of Figure 2a, given the training chirp stimulus protocol. From the example, GNE can capture firing dynamics better, especially within the green rectangle. We further quantify the fitness of the predicted model responses using MAE and Pearson correlation coefficient between predicted model responses and the groundtruth. In this case, GNE outperformed MOO and CNN with 1.8 and 4.1 times performance gain in terms of MAE and the predictions from GNE have 2.1 and 2.2 times higher Pearson correlation coefficient than MOO and CNN correspondingly (Figure S2a).

We also evaluate the noise robustness of three algorithms by applying Gaussian noise at different SNR levels to the model responses in the test phase (Figure 2c), where the noise is added to the neural response, and then the noisy response serves as the input to the deep neural network to generate parameter prediction. Then, the predicted parameter and the same stimulus protocol during training are applied to the neuronal model to obtain the predicted responses. We find that MOO is quite stable to Gaussian noise, although its performance is not quite good. GNE showed an obvious trend of error decreasing as the SNR went higher and the overall performance is several orders superior to MOO, and the parameters’ predictions of GNE are more accurate than that of CNN at all SNR levels.

We also evaluated the impact of training scale on the prediction accuracy (Figure S2b). As the training scale increases, the prediction error of parameters decreases significantly. The prediction accuracy of GNE given 100K training data is comparable to that of CNN with 1M data (Figure S2b).

Although the model response predicted by MOO and CNN cannot fully replicate the firing patterns, they can at least replicate the burst firing patterns when the amplitude of stimulus gets larger (Figure 2b & c). To further test which kind of firing patterns can be captured under the given the predicted parameters of MOO, CNN, and GNE, we evaluated the model generalization ability by applying stimulus protocols which are not presented during the training or search phase. We adopted stimulus protocols from (Podlaski et al., 2017), with modifications on the amplitude and base values of the stimulus protocols (Figure S3). We use MAE and Pearson correlation coefficient as the metrics to quantify the model generalization ability under 42 stimulus protocols with differences in amplitude and shape. GNE is superior to both in all cases except one with the lowest stimulus amplitude (Figure S3a).

Compared with MOO and CNN, the performance boost demonstrates that the features extracted by GNE is more powerfully, has stronger robustness towards noise and can generalize better to unseen stimulus protocols than the hand-crafted features and features extracted by CNN. Therefore, we analyzed why GNE outperformed CNN from the perspective of features. We first plotted the correlation heatmap among features extracted from GNE and the baseline both before fully connected layers (Figure 2d, e). From the correlation heatmap of GNE (left of Figure 2d), we can see a cluster of sparsely correlated features labeled as 1 and strongly correlated features labeled as 2, and the whole landscape of features is shown (right of Figure 2d). The color of the features corresponds to the labels in the heatmap. The features in the first cluster include those describing firing patterns of bursts of spikes and the single spike. While the second cluster mainly contains features describing irregular and dynamical fluctuations. Features extracted from CNN architecture show a different pattern and can be divided into 3 clusters (left of Figure 2e). The first two clusters are all describing firing patterns of bursts of spikes, while in the third cluster, only few features are encoding single-spike level features, and there are no features encoding subthreshold fluctuations (right of Figure 2e). With features from different levels of neural responses, GNE outperformed the baseline in all the four neuronal models in this paper. It can be inferred that features describing subthreshold dynamics are critical determinants to constrain the parameters. Therefore, lack of features describing subthreshold dynamics may account for the poor performance of hand-crafted features in MOO and the features depicting firing patterns in CNN.

### Inferring Single Neuronal Models on Experimental Recordings

All the above evaluations demonstrate that GNE outperforms MOO and CNN on synthesized data, while in this section we show that GNE also can be applied to experimental recordings. In this particular case, we use the same HH single-compartment model as illustrated in equation 7 and 8. There are 8 free parameters to predict, which are *g*_Na_, *g*_K_, *g*_L_, *g*_M_, *τ*_Max_, *d*_T_, *σ*, *E*_I_. The lower bound of them is [0.5, 10−4, 10^−4^, 10^−4^, 50,−90, 10^−4^,−100] and the upper bound is [80, 15, 0.6, 0.6, 3000,−40, 0.15,−35]. We trained the deep neural network on 800K synthesized data (see methods for details of the training configurations and dataset generation) until the validation loss converged, and the well-trained estimator is loaded with experimental recordings as the input to output the corresponding model parameters (Figure 3a). We demonstrate the prediction results of experimental recordings of mouse spiny neurons from Allen Institute(Allen Cell Types Database, 2016) (Figure 3b; for results of other 18 neurons see Figure S12a). We can see that from a single pass of prediction, GNE can precisely replicate a wide range of experimental recordings (the first 5 examples in Figure 3b, Figure S12a). Some experimental recordings that cannot be correctly estimated through single training pass (the last example in Figure 3b). Now, we’ll show that how to combine GNE with SNPE-based algorithms to calibrate a specific experimental recording. SNPE-based inference algorithms shed lights on accurately estimating parameters of neuronal models, as proposed in (Gonçalves et al., 2020). Even though, most of the applications still need lots of efforts for manual feature extractions, which is non-trivial (Figure 3d). GNE can extract temporal dynamics from raw neural responses. On account of it, we present that GNE can serve as the feature extractor for SNPE algorithms to get better results, where the fully-connected components in our estimator are replaced by SNPE component (Figure 3c). We compared the predicted results with SNPE whose inputs are manually crafted features (Figure 3d). The marginal density distributions of maximal conductance variables are illustrated in Figure 3c and d. From the density map, we can clearly see that the variance of conductance of ion channels from the combination is much smaller than that from SNPE alone (Figure 3e), which suggests that the temporal features extracted from GNE can better constrain model parameters because it efficiently captures temporal dynamics. The prediction from GNE plus SNPE can better replicate the dynamics as observed in experiments than SNPE with hand-crafted features (Figure 3f).

**Figure 3:**
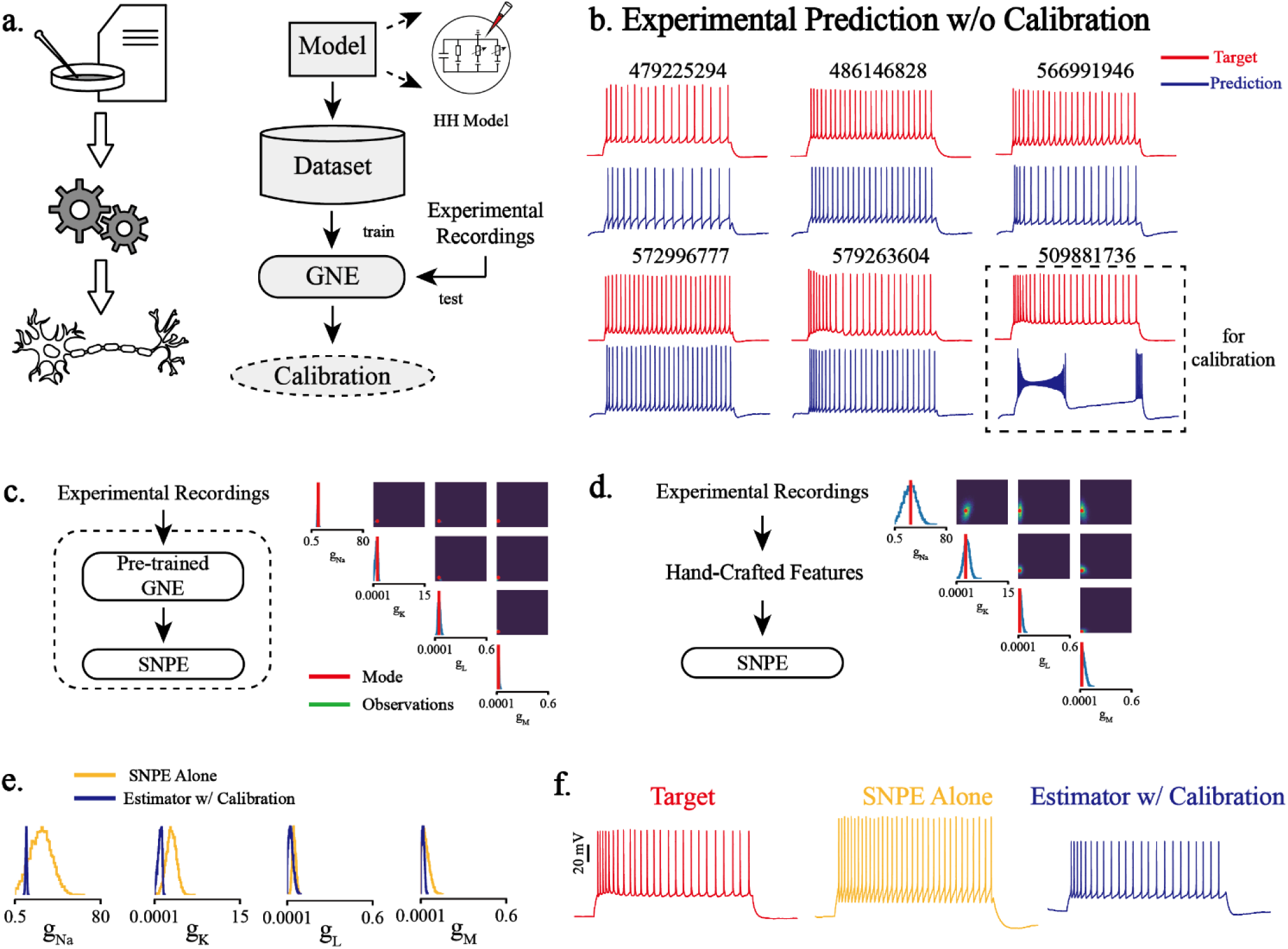
Results to predict model parameters from experimental recordings. **a** The roadmap and method workflow to estimator model parameters from experimental recordings. First synthesized dataset generated from computational models are used to train the estimator until training loss converges. Then experimental recordings serve as the input to the estimator to get the corresponding model parameters. If the direct prediction of parameters is not accurate enough, statistical calibration is applied to fine-tune on specific experimental recordings. **b** 6 examples from the Allen Institute Cell Types Database, the upper numbers represent the cell specimen ID in the database. The last one is the failed prediction results from the single training pass of an experimental recording from Allen Institute Cell Types Database, which will be statistically calibrated further. **c** The method workflow and results to use statistical calibration on specific experimental recordings with SNPE alone, where the input for the combination is the hand-crafted features of raw experimental recordings. (Only maximal conductance variables are shown, for full results see Supplementary) **d** The method workflow and results to use statistical calibration on specific experimental recordings with the combination of estimator and SNPE, where the input for the combination is the raw experimental recordings. **e** A comparison of the estimated marginal distribution density map of maximal conductance variables between SNPE alone and estimator with SNPE. The variance of the density of estimator with SNPE is much narrower, which denotes a higher confidence. **f** The result of calibration the failure case shown in **b**, the left is the experimental recordings, the middle and the right are generated using the mode from SNPE with hand-crafted features, and combinations of estimator and SNPE.

### Inferring Connectivity Conductance of a Small Biophysical Neural Microcircuit

The above sections discussed the accuracy, generalization ability and noise robustness of GNE in the optimization of single neuronal models. In this section, we show that GNE can accurately infer the dynamics of neural circuits based on well-constructed neuronal models (Figure 4a). We built a neural circuit upon 3 precisely constructed HH L5 pyramidal neurons in the mouse primary visual cortex rom Allen Institute Cell Type Database(Allen Cell Types Database, 2016) (Figure 4b, see Methods for details). Each connection is composed of a NMDAR and an AMPAR (Figure 4b). We used GNE to estimate the conductance of each receptor in the circuit (12 parameters in total). In this case, each neuron receives a clamped square current with Gaussian white noise. The prediction MAE of the synaptic conductance is 0.068 and 0.025 for NMDAR and AMAPR correspondingly (Figure 4c). We demonstrate the predicted NMDA and AMPA current when two spikes are transmitted from the pre-synaptic neuron in Figure 4d. Both predicted currents can precisely reproduce the groundtruth. This application proposes insights for further neural circuitry reconstruction. For precise reconstruction, it can be divided into two parts, namely, first to reconstruct single neurons and then reconstruct neural circuits built upon those neuronal models.

**Figure 4:**
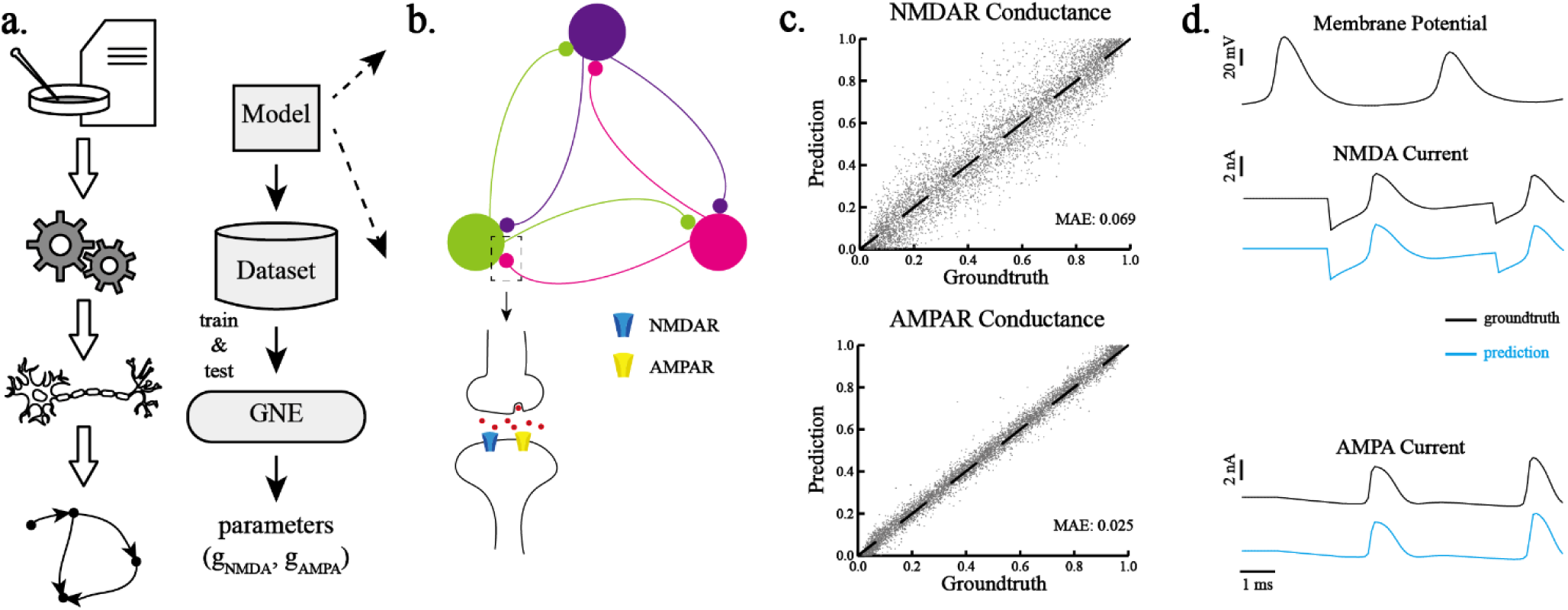
Results to estimate the synaptic weights of a neural microcircuit with already-reconstructed single neuronal models. **a** The roadmap and method workflow to reconstruct neural circuitry with single neuron reconstruction as the cellular bases. **b** An illustration of the neural microcircuit composed of 3 HH neurons is shown, where neurons are fully connected with each and the self-connections are eliminated. Each connection is composed of a NMDAR and an AMPAR. The input to each neuron is clamped. The responses of the three neurons together serve as the inputs to the estimator, and the output are the connectivity weights among the six neurons. **c** Prediction accuracy of the conductance for NMDAR and AMPAR. **d** The comparison of the predicted NMDA current and AMPA current with the groundtruth when two spikes are transmitted from the pre-synaptic neuron.

Further we compared the performance of GNE with I&F method (Ladenbauer et al., 2019) on a neural circuit composed of 6 fully-connected I&F neurons. There are total 30 synaptic weights to estimate and the ranges of the weights are all within [-0.75, 0.75]. Limited by experiments recordings, we can only obtain the spike timing of each neuron in the neural microcircuit, so we first use the firing rates calculated from the spike train as the input of GNE (for details, see Methods). In this scenario, our prediction accuracy is 1.6-2.7 times higher than I&F method (Figure S14a). As subthreshold signals contain richer information than spike train, the accuracy with subthreshold as the input is 2.0-3.6 times higher than using only spike train as the input (Figure S14a). In both cases, our prediction can nearly perfect match the target dynamics regardless of the input as spike train or subthreshold dynamics (Figure S14b). Given the discoveries that the prediction accuracy is higher with subthreshold dynamics as the input than with spike train, it suggests that subthreshold dynamics within a neural circuit might be of great importance in terms of its dynamics or functions.

### Inferring Connectivity of a Large-Scale Neural Circuit

In the previous section, we showed prediction results of the connectivity weights of a neural circuit with clamped. We used the responses of all neurons in the circuit to predict all connectivity weights at the same time. This kind of approach is only suitable for circuits of a small size which will face the problem of scalability when the size of the circuit grows large. Also, the stimulus inputs to neurons in the circuit is clamped and such experimental techniques may be infeasible when dealing with a large neural network. Therefore, in this section we present a training scheme (Figure 5a) to solve the scalability to the size of neural circuits and experimental recordings, to deal under conditions with background noise of high variation. In this case, we constructed a fully-connected neural circuit with 100 I&F neurons. When generating the training dataset, for each data entry, we simulate the neural circuit for 20 times, each time with a different background condition (see Methods for details). To solve the scalability to the size of the neural circuit, we choose to estimate the connectivity weights pair by pair. As shown in Figure 5a, the inputs to GNE are the responses of two neurons within the circuit each time, and the outputs are the connectivity weights between the two neurons. To deal with arbitrary number of experimental recordings, we fuse the feature representation after Res-LSTM module before further processed by the fully-connected units. We visualized the predicted weight matrix and its associated reconstructed neural circuit in Figure 5c, and our prediction and reconstruction can almost recover the properties of the groundtruth. The recordings of each neuron in Figure 5b are shown in Figure 5c, where we can see the firing patterns under different recordings are diverse. Features extracted from recordings are fused by performing average operation. Feature representation of singe recording contains sparse information in all scenarios (excitatory, inhibitory and no connections, Figure 5d). By fusing feature representations under different background conditions, the fused feature representations have richer and more concrete information to decide the type and the weight of each connection (Figure 5d). In this way, the method is both scalable to the size of the circuit and the number of experimental recordings. Our algorithm can still give more accurate prediction compared with I&F method (Ladenbauer et al., 2019) and GLMCC method (Kobayashi et al., 2019) (Figure 5e). Our algorithm benefits more as the number of experimental recordings increases (Figure 5e). Moreover, we found that there is a clear imbalanced preference to identify synaptic connections as inhibitory ones for both I&F method and GLMCC method (Figure 5e). GNE doesn’t encounter the prejudice for inhibitory connections and performed similar and high identification accuracy for both excitatory and inhibitory connections.

**Figure 5:**
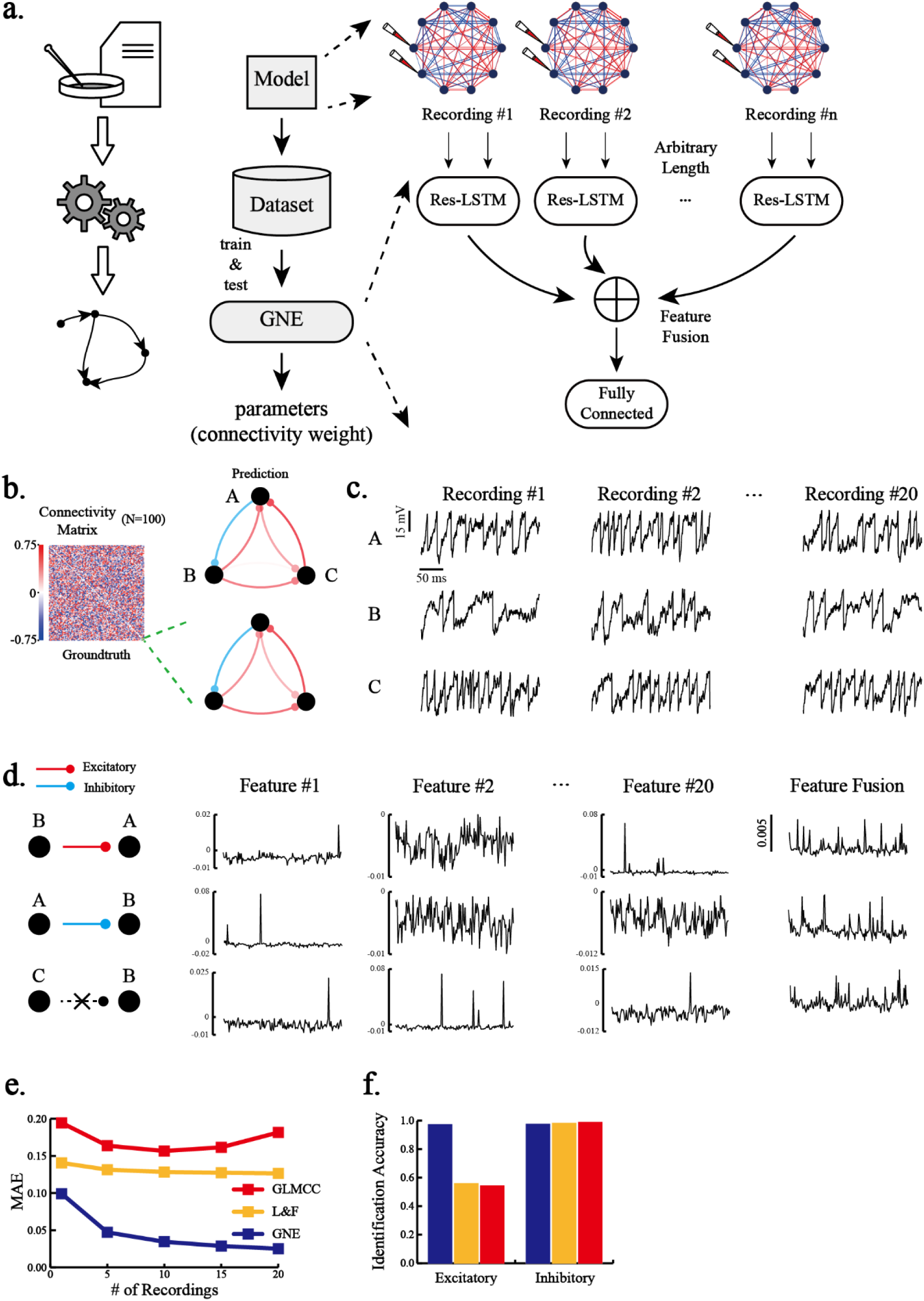
Results to estimate the synaptic weights of a neural circuit with random inputs. **a** The roadmap and method workflow to estimate synaptic weights of a neural circuit with random inputs. The dataset is generated by simulating the neural circuit for n times, where the background input each time is different. The training scheme to deal with arbitrary number of experimental recordings. Responses of two neurons in the neural circuit serve as the input to the estimator and features extracted by the Res-LSTM are averaged before they are finally regressed by fully connected units. **b** The left matrix is the groundtruth connectivity matrix of the whole neural circuit with 100 neurons. The middle column shows the predicted and groundtruth connectivity matrix of the sub-circuit within the dashed box and the right column shows the corresponding circuit reconstruction visualization, where blue lines are negative connections and red lines are positive connections. The shade of the color represents the strength of the connection. **c** Neural responses of the three neurons in **b**. Three recordings are selected for each neuron, the time length illustrated in the figure is 0.25ms, while the length for training is 2.5s. **d** The illustration of the effect of feature fusion in **a**. The features extracted from single recording and fused among 20 recordings are shown, where we can see the information in the last is denser to reflect the connection of the pair. **e** The relationship between prediction accuracy and the number of experimental recordings. As the number of experimental recordings increases, the prediction MAE of both algorithms decreases. The accuracy of our algorithm is higher in all cases. The unit time for a single recording is set as 2.5s for GNE and I&F method and as 5s for GLMCC method. **f** Identification accuracy comparison for excitatory and inhibitory connections. GNE can achieve balanced and high identification accuracy for both types of connections, but L&F and GLMCC method show a clear prejudice for inhibitory connections and perform poorly on excitatory connections.

## Discussion

Experimental neuroscience and computational neuroscience are promoting the development of each other from different perspectives, where the bridge between the sub-fields of neuroscience lies in the gap between experimental discoveries and model discoveries(Abbott, 2008; Q. J. M. Huys et al., 2016; Wang et al., 2020). With the development of experimental techniques, the observation granularity goes finer and finer, from dendritic morphologies to molecular pathways. As a result, the model design should be aligned with the experimental discoveries and more compartments are considered in a single detailed neuronal model and neural circuit models(Larkum et al., 1999; Du et al., 2017; Ebner et al., 2019; Hjorth et al., 2020). However, the massive number of free parameters in the neural model makes it hardly possible to fully replicate experimental discoveries in a complex neural model. Until now, the mainstream approaches still rely heavily on carefully hand-crafted features for different neuronal models and various neural responses. The intense feature engineering is non-trivial and extraordinary time-consuming, which greatly hinders the further development of neuroscience. Here in this paper, we propose GNE, an LSTM-based deep learning method to infer the parameters of single neuronal models and connectivity properties of a neural circuit, and a roadmap based on it to reconstruct single neurons and neural circuits. The method is equipped with a powerful and robust feature extractor, which can efficiently capture temporal dynamics of neural responses. GNE can lead to accurate parameter predictions for a wide range of single neuronal models or neural circuits, and it is robust to noise interference, and can well generalize to other unseen stimulus protocols. In summary, the reconstruction roadmap and GNE might shed light on further experimental design, computational model design and brain reconstruction research.

### Related Work

Researchers have spared great efforts in designing parameter optimization approaches for decades(Prinz et al., 2003; Schneider et al., 2004; Keren et al., 2005; Gerken et al., 2006; Druckmann et al., 2007; Gurkiewicz & Korngreen, 2007; Menon et al., 2009; Hendrickson et al., 2011; Rossant et al., 2011; Santana et al., 2011; Bahl et al., 2012; Svensson et al., 2012; Markram et al., 2015; Eliasmith et al., 2016; Rumbell et al., 2016; Masoli et al., 2017; Neymotin et al., 2017; Gouwens et al., 2018; Kobayashi et al., 2019; Ladenbauer et al., 2019; Billeh et al., 2020; Endo et al., 2020; Gonçalves et al., 2020).

As for algorithms designed for single neurons, conventional approaches can be roughly divided into model-agnostic approaches which aim at a unified solution for any kinds of neuronal models(Prinz et al., 2003; Schneider et al., 2004; Keren et al., 2005; Gerken et al., 2006; Druckmann et al., 2007; Gurkiewicz & Korngreen, 2007; Menon et al., 2009; Hendrickson et al., 2011; Rossant et al., 2011; Santana et al., 2011; Bahl et al., 2012; Svensson et al., 2012; Eliasmith et al., 2016; Rumbell et al., 2016; Masoli et al., 2017; Neymotin et al., 2017; Gouwens et al., 2018) and model-specific approaches which aim at a precise or guaranteed-optimal tuning for certain types of neuronal models(Bush et al., 2005; Q. J. Huys et al., 2006; Q. J. Huys & Paninski, 2009; Pozzorini et al., 2015). Model-agnostic approaches, in particular evolutionary or genetic algorithms, are among those popular methods. These approaches are formulated as a single- or multi-objective optimization (MOO) procedure, which search sets of parameters that can minimize the errors of pre-defined objective functions based on pre-defined features. Possible weaknesses of these kind of methods might lie in that they normally rely on hand-crafted features and well-designed loss functions and massive manual interventions are generally required. As for model-specific approaches, typical examples include variations of linear regression(Bush et al., 2005; Q. J. Huys et al., 2006), expectation-maximization algorithms(Q. J. Huys & Paninski, 2009), convex optimization(Pozzorini et al., 2015). They appear to be more effective to certain types of neuronal models with similar mathematical representations, but lack of generality on other types of neuronal models. All these traditional methods have made significant contributions to the computational modeling in the field, however, their limitations mentioned above have greatly weakened applications of the traditional methods on more complicated or large-scale models, such as biophysically detailed models or large-scale circuit models. It was only recently that deep neural networks (DNNs) have been formally introduced into neuronal model optimizations, which shed lights on parameter-tuning problems. Deep neural network, also called “Deep learning”, is perhaps the most popular technique in the modern AI and has reshaped the machine learning community during the last decade(Krizhevsky et al., 2012; Sutskever et al., 2014; He et al., 2016; Yin et al., 2016; Gamboa, 2017; Brown et al., 2020). Deep artificial neural networks have the following interesting properties: (1) they are powerful feature extractors;(2) in theory, they are guaranteed to approximate any mathematical functions(Leshno et al., 1993), which makes it an ideal tool to “approximate” mathematical models underlying neural activities. (Ben-Shalom et al., 2019) first applied a 5-layer convolutional neural networks (CNNs) to optimize conductance-based neuronal models, which can directly incorporate raw data of neuronal dynamics and output the parameters for any models. This approach is very straightforward and was shown to outperform the conventional MOO method on a variety of neuronal models. As a variation of deep learning approaches, an advanced Bayesian-based approach(Gonçalves et al., 2020), developing upon SNPE(Papamakarios & Murray, 2016; Lueckmann et al., 2017; David et al., 2019) and autoregressive flow(Papamakarios et al., 2017), also came out lately. Instead of directly output the free parameters of models, it estimates the posterior distribution P(θ|x) of the free parameters θ from summary features x, which endows more flexibilities and interpretability in parameter tuning. However, one potential weakness in either CNN-based or Bayesian-based approaches is likely to be their feature extraction process: (1) To the CNN-based approach, it employs single CNN as feature extractors, which treat the neural responses at each time step as independent events. In deep learning community, it is well established that CNN architecture alone cannot well perform in temporal-related tasks without special design(Lea et al., 2016; Yin et al., 2016), such as natural language processing. Thus, features extracted from CNN might lose important temporal details. (2) To the Bayesian-based approaches, they critically rely on carefully hand-crafted feature engineering and the related hyper-parameters(Gonçalves et al., 2020), which might limit its applications to those neural dynamics without knowing sufficient prior knowledge.

As for algorithms designed for neural circuits, search-based algorithms(Markram et al., 2015; Billeh et al., 2020) and SNPE-based algorithms(Gonçalves et al., 2020) can still be applied, while they also requires non-trivial feature extraction currently. (Ladenbauer et al., 2019) tries to reconstruct neural circuits by fitting experimental recordings to I&F neural models. By performing likelihood maximization, the connectivity weights of the neural circuit can be estimated. (Kobayashi et al., 2019; Endo et al., 2020) estimate neural connectivity through the cross-correlogram(CC) from the spike trains of two neurons. (Kobayashi et al., 2019) fits the CC through a generalized linear model(GLM), and (Endo et al., 2020) estimates neural connectivity from CC by training a CNN. While the above three approaches can only estimate a rough synaptic weight and the existence of the connection, which they cannot estimate the connectivity strength of multiple synaptic receptors (NMDAR and AMPAR for example).

### Subthreshold Dynamics Might be Critical to Constrain Model Parameters

As is shown in Results, GNE can extract more diverse sets of features. Both GNE and CNN architecture can efficiently capture massive features regarding consecutive spikes. More than this, GNE comes to extract more features describing single spike information and irregular subthreshold fluctuations (Figure 2d). We can interpret these features as precise spike timing and the relationship between two spikes within a long-time window and important information regarding subthreshold dynamics. Also, we achieved high estimation accuracy in estimating synaptic weights by using subthreshold dynamics as the input than using spike trains (Figure S14b).

Those phenomena might indicate that features depicting single spikes and subthreshold dynamics should be critical in constraining a neural model or even its functional role. However, in previously presented algorithms(Druckmann et al., 2007; Rumbell et al., 2016; Masoli et al., 2017; Gouwens et al., 2018), a majority of hand-crafted chosen features only focus on only firing patterns, such as first spike timing, firing rates, and et al.. Subthreshold dynamics are considered as a representation of background and input activity statistics(Clay & Shrier, 1999; Hillenbrand, 2002). Subthreshold dynamics have been pointed out to be important for the neural system in terms of firing pattern association(Otomo et al., 2020), information integration(Ratté et al., 2015), plasticity(Latorre et al., 2016), and they also play an important role in neural networks and neural circuits(Ness et al., 2016; Wright & Wessel, 2017). The discoveries based on GNE also showed that subthreshold dynamics are extremely important in identifying neuronal models, which might open a door for further investigation about the functional role of subthreshold dynamics for single neurons or neural circuits. The discoveries from our extracted features might also cast light on hand-crafted features and some works have already focused on spike prediction based on subthreshold dynamics(Kobayashi & Shinomoto, 2007).

### GNE is Efficient in Utilizing Big Data and Can be Easily Applied

As it is shown in our evaluations, deep learning showed its magnificent ability in processing and utilizing big data. Theoretically, we can generate an infinite number of data from a prior parameter distribution and a computational model, so that we can improve the performance of deep learning in terms of training size (shown in Figure S2d, Figure S4d, Figure S6d, Figure S8d, Figure S10d). Search-based approaches, such as MOO, can only estimate parameters by resampling and re-searching combinations of parameters independently, which means the set of parameters sampled for one experimental recording cannot be used for future unseen ones. Its inability to use big data is a huge time consumption. On the other hand, SNPE-based approaches shed light on solving this problem by utilizing amortized learning(Speiser et al., 2017; Webb et al., 2018), which can estimate the posterior distribution for all observations within the parameter space. Even though, they still require fine-tuning on each observation by resampling and re-training to obtain accurate predictions. In contrast, as for GNE, once a deep learning model is trained, it can be used for different experimental recordings directly without further modifications. Therefore, besides accuracy, the time consumption of deep learning is less and GNE is more scalable than search-based algorithms. The re-usage of the trained model allows neuroscientists to evaluate their models much more quickly. As for GNE, one can predictthe parameters within 5 seconds by loading the pre-trained models on an NVIDIA V100 GPU. In addition, GNE is straightforward and highly user-friendly for neuroscientists, who only need to provide a target neural response and a training dataset, without any necessities to modify the deep learning architecture or the rawresponses.

### GNE can be Directly Applied to Experimental Data

GNE can well predict dynamics of experimental recordings (Figure 3b). However, it is still worth noticing that it is not a panacea for all experimental recordings. We tried other experimental recordings of mouse spiny neurons from Allen Cell Types Database(Allen Cell Types Database, 2016), and found that GNE cannot predict some of them well. Two main reasons may account for this: 1. the HH single-compartment neuron model cannot mathematically or theoretically replicate the experimental recordings, which means there is no set of parameters can that produce the same experimental data to perfection. 2. GNE is highly effective on extracting features of synthesized data, but it cannot guarantee that those features can present in the experimental recordings. Naturally, the features of the experimental data may differ from those of synthesized data, and the differences may account for the imperfection prediction. The first problem cannot be well tackled through the modifications of optimization algorithms, while the second one can. Just because of this, we present the synergy of pre-trained Res-LSTM and SNPE to calibrate on a specific experimental recording (Figure 3c).

### GNE Provides Statistical Calibration on Experimental Recordings

Since now, two dominant approaches in constraining neuronal model parameters, search-based approaches and Bayesian-based approaches, are still rely heavily on carefully designed features, which is non-trivial. Here, GNE provides a powerful and robust feature extractor that can capture temporal features efficiently and automatically. Because of this, GNE does not conflict with the above two methods, and one can use the proposed pre-trained models as the front-end of those methods to benefit from our extracted features. The combination of GNE and the above two approaches will finally unite the advantages of each approach in three aspects. First of all, it can efficiently extract features, which can capture not only the sub-threshold temporal dynamics, but also the spike timing features. Second, it can generate a set of feasible parameters rather than a single parameter estimation, since different combinations of parameters may provide similar model responses(Goaillard et al., 2009; Gouwens et al., 2018; Gonçalves et al., 2020). Third, it will be equipped with interpretable analysis to inspire further model design(Gonçalves et al., 2020). We showed that the prediction by the combination of the two methods is more accurate than SNPE alone (Figure 3f). This allows for calibration on specific experimental data to get more accurate predictions.

### GNE might Inspire Future Experiment Design on Brain Reconstruction

Brain reconstruction has been a hot field in recent years, from the brain of drosophila(Mann et al., 2017; Franconville et al., 2018) to rats(Markram et al., 2015; Billeh et al., 2020). Those research works aim at building a reconstructed digital neural system of the brain of the above species, from functional to biophysical scale. However, due to the limitations of experimental techniques and reconstruction methods (i.e. parameter estimation algorithms), the reconstruction at synaptic level is majorly based on statistical information which leads to a coarse reconstruction of synaptic properties. GNE sheds light on the precise reconstruction of synapses at the network level, which is both scalable to the size of the neural circuits and the length of recording time. First, GNE can give precise single neuron reconstruction within a neural circuit. Then, by multi-site intracellular or extracellular recordings, or calcium fluorescence imaging techniques, together with biophysical neural circuit mathematical modelling, GNE can give connectivity or synaptic reconstruction of the circuits. Such precise reconstruction from single neuron to neural circuit might inspire the future design of experiments as well as computational modelling.

### Criteria on Judging the Fitness of Neuronal Model Might Have Bias and Should be Carefully Designed

There have been many efforts to design reasonable criteria to judge the fitness of a model on experimental recordings(Weaver & Wearne, 2006; Druckmann et al., 2008; Jolivet, Kobayashi, et al., 2008; Jolivet, Roth, et al., 2008; Gerstner & Naud, 2009; Schrimpf et al., 2018; Schrimpf et al., 2020). However, a consensus hasn’t been reached. Here, in this paper, we selected two categories of evaluation metrics, one for predicted parameters and one for predicted model responses.

We also analyzed the two response prediction criteria, curve MAE, and Pearson correlation coefficient, on all the neuronal models studied and found that the two criteria have a bias on the neural firing frequency. Curve MAE suggests point-to-point level similarity, while the Pearson correlation coefficient strengthens the linear association between two responses. A criterion shall maintain stable under different evaluation conditions, such as different stimulus amplitudes. Here we analyzed how the two criteria behaved when giving step stimulus protocols to neuronal model at different amplitudes. We demonstrated the scores given by the two criteria versus neural firing frequency (Figure S15). As for curve MAE, we found that all these methods tended to increase and then decrease on all the neuronal models studied (Figure S15). This phenomenon suggests that the stimulus protocol used to evaluate a computational model is critical because even when the model is given a great score at an extremely high firing frequency. It is quite likely that the frequency where the criteria show the extremum can be a base condition for neuroscientists to judge their computational models as well as the associated parameters. It also provides insights into the design of proper stimulus protocols experimentally to fully discriminate neurons within a large population.

On the other hand, parameter estimation algorithms usually do not emphasize the importance of generalization to unseen stimulus protocols. In (Gouwens et al., 2018), they discovered that many parameters combinations showed high accuracy under the same stimulus protocol during searching. When an unseen stimulus protocol was applied to the model, these parameter combinations performed poorly. Therefore, the generalization to unseen stimulus protocols should be a crucial criterion to evaluate the performance of the predicted parameters. In this paper, we tested our model generalization ability to a wide range of stimulus protocols, and GNE outperformed MOO and CNN.

In summary, the criteria for judging the fitness of a neuronal model and its associated parameters should be carefully designed, and all aspects that may cause bias under various conditions should be considered. A systematic and thorough study and design of such criteria remain future research and discussion.

## Data Availability Statement

The code of our method will be published online upon acceptance.

## Methods

### Configurations of Computational Models

#### Stimulus Protocols

For single neuronal model on synthesized data evaluation tasks, we used the chirp-like stimulus protocols to generate the training dataset, which are widely used in experimental neuroscience to study phase lock or invariant neuronal responses, especially in the auditory system and retina(Petoe et al., 2010; Baden et al., 2011; Pushpalatha & Konadath, 2016; Schröder et al., 2020; Szatko et al., 2020). As for single neuron inference on experimental data, we use step current with Gaussian white noise. As for neural circuit models, we use Gaussian current stimulus protocols. The details of stimulus protocols can be seen in the following.

#### HH Single-Compartment Neuron Model

The HH single-compartment neuron model is defined as equation (7)-(8). The stimulus duration is 200 ms with 0.5 ms as the time step. The protocol for these two models is as the following:

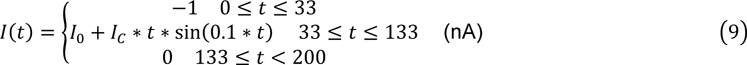

where *I*_0_ is chosen as 0.015, *I*_*C*_ is chosen as 4.

#### Izhikevich Neuronal Model

The Izhikevich neuronal model we used is the standard vision presented in (Izhikevich, 2003). The computational mechanisms are as the following:

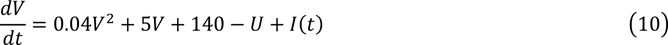

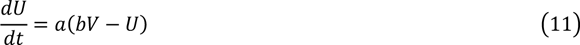

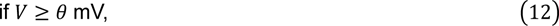

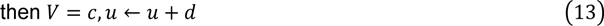

where θ is the firing threshold which we set as 35 mV, and *a*, *b*, *c*, *d* are the free parameters. The stimulus duration is 200 ms with 0.5 ms as the time step. The protocol for these two models is as equation (11), where *I*_0_ is chosen as 0.05, *I*_*C*_ is chosen as 10.

#### HH Ball-Stick Model

We created a two-compartment ball-stick HH model in NEURON simulator(Carnevale & Hines, 2006), a soma and a dendrite. The diameter of the soma is 21 microns. The diameter of the dendrite is 1 micron and length of it is 100 microns. There are four active ion channels, namely Na, K_m_,Ca, K_Ca_, and a passive leaky channel on the dendrite. Soma has one more K_v_ ion channel than the dendrite. The maximum conductance of K_Ca_ channel on both the soma and the dendrite is fixed to 10 pS/*μ*m^2^. The maximum conductance of K_v_ channel on the soma is fixed to 5 pS/*μ*m^2^. The axial resistance is set to be 100 *Ώ* ⋅ cm. The membrane capacity is set to be 1 mF/cm^2^.

#### Mainen-Sejnowski Model

We use the NEURON simulator(Carnevale & Hines, 2006) to simulate neuronal model responses in this model. The source code of the model is obtained from ModelDB(McDougal et al., 2017) at http://modeldb.yale.edu/2488. We chose L4 stellate neuron and L5 pyramidal neuron to perform the experiments without any modifications on the default parameters except the free parameters to be estimated. The stimulus duration is 500 ms with 0.1 ms as the time step. The protocol for these two models is as the following:

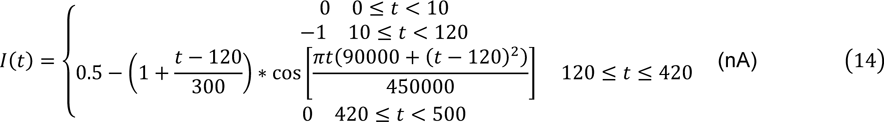

#### LIF Neural Circuit Model

We use the code published by(Ladenbauer et al., 2019) at https://github.com/neuromethods/inference-for-integrate-and-fire-models to simulate the neural microcircuit. For the evaluation with spike train as the input of GNE, we transform the binary spike train into a consecutive firing rate curve. We calculate the average firing rate of each neuron within the time window of 5 ms. The amplitude of the background activity to each neuron has mean value of 1.4 mV/ms and standard deviation of 0.7, and the standard deviation of the background activity to each neuron has mean value of 2 and standard deviation of 1.

#### HH Neural Microcircuit Model

We use three reconstructed neurons from Allen Institute Cell Types Database(Allen Cell Types Database, 2016) whose IDs and sweep IDs are (566991946, 36), (572996777, 32), (596282602, 38). All the HH equations are the same as equations (7) - (8), and the corresponding free parameters the direct estimation from GNE as mentioned in Figure 3. The mean of inputs to each neuron are randomly chosen within [0.9, 1.4] nA and then clamped, with Gaussian white noise of standard deviation 0.05 nA. The neural microcircuit is fully connected, there are an AMPA receptor and a NMDA receptor for each connection. The AMPA-mediated current is adopted from (Tsodyks & Markram, 1997; Kobayashi et al., 2019). The NMDA-meditated current is adopted from (Destexhe et al., 1998; Kobayashi et al., 2019).

### Training Configurations

After acquiring a large amount of training data, it is possible to train the deep neural network using the backpropagation algorithm(Rumelhart et al., 1985). The pipeline of training is illustrated in Figure 1a. For four single neuronal models and LIF neural circuit models with clamped inputs, we divided the training dataset into training data and validation data at the ratio of 9:1. The default size of the training dataset is 1M. For the LIF neural circuit models with random inputs and HH neural circuits models, we set the training to validation ratio to 99:1. The training size of HH neural circuit models with clamped inputs is 5M. The training size of the LIF neural circuit models and HH neural circuit models with random inputs is 1K 100-fully-connected neuron circuits. Then we sampled another 10K amount of data as the test dataset. We normalized the parameters of neuronal models to [0,1] for GNE and [−1,1] for CNN. We used a batch size of 200 and chose Mean Squared Error(MSE) as the loss function to calculate the error between the prediction and the groundtruth of the parameters for both GNE and CNN. We set the initial learning rate as 0.0001 and used Adam optimizer(Kingma & Ba, 2014). This procedure is repeated iteratively until the validation loss doesn’t decrease for more than 20 epochs or reach a total of 100 epochs. For the evaluation of the MAE of predicted parameters and the groundtruth, we transformed the normalization to [0,1] for CNN.

Compared with the original ResNet-50 design, we modified the first input channel from 3 to the number of model response sequences, and replace the ReLU activation function with the Leaky ReLU activation function. We also eliminated the average pooling layer at the bottom of the original ResNet-50, and attached a 5-layer fully connected layers to our implementation. The number of units in each fully connected layer is [512,256,256,128,128] correspondingly.

### Deep Learning Baseline and Multi-Objective Optimization Configurations

(Ben-Shalom et al., 2019) is the first one to use deep learning to infer parameters of conductance-based neuronal models at present, so we use its architecture as our deep learning baseline. We name this architecture as CNN in the following sections and figures. For the sake of convenience and simplicity, we only trained one deep learning model for each computational model, instead of 32 independentones, andwe use therawneuronalmodel responses as the input without data transformations as presented in (Ben-Shalom et al., 2019). We also compare our deep learning architecture with MOO algorithms. We use (µ + *λ*) as our evolutionary strategy and NSGA-II selection operator to select the best parameters from the population. We initialize the algorithm with 1000 individuals as the population and generate 1000 off-springs at each generation with 0.7 as the crossing probability and 0.3 as the mutation probability. The iteration stops when there is no improvement for 10 epochs or the overall number of individuals generated exceed 1 million.

### L&F Method

The L&F is proposed by (Ladenbauer et al., 2019) and we used the published code at https://github.com/neuromethods/inference-for-integrate-and-fire-models to infer the synaptic weights of L&F neural circuits. We used the original sets of hyperparameters in the code without any modifications.

### GLMCC Method

GLMCC method is proposed by (Kobayashi et al., 2019) and we used the published code at https://github.com/NII-Kobayashi. We set the membrane time constant as 20 ms, and synaptic delay as 1 ms, which are the same as the L&F neurons used for simulation. The time length for cross-correlogram is 100 ms with 1 ms time bin. Other hyperparameters are remained unchanged as in the original codes. After estimation, we got the predicted weights *w*. In order to recover the true weights for performance comparison, we scaled the predicted weights to α*w*, where the α is obtained by performing simulated annealing algorithm to minimize the MAE between *αw* and true synaptic weights based on results from 20 recordings.

### Evaluation Metrics

There is no recognized metric to judge how well a neuronal model together with its free parameters matches experimental recordings. To eliminate metric bias when making comparisons, we used three different quantitative metrics to compare GNE, the deep learning baseline, and MOO, namely:

1. Mean Absolute Error (MAE) of predicted parameters and groundtruth parameters
2. Pearson Correlation Coefficient between the model responses generated from predicted parameters and groundtruth model responses
3. MAE between the model responses generated from predicted parameters and groundtruth model responses

In addition to applying different evaluation metrics, we also implemented two streams of evaluation tasks.

#### Noise Robustness Evaluation

Experimental recordings are naturally noisy, therefore even if there is no noise in the training phase, the parameter estimation algorithm must function normally under the noise condition. Hence, we evaluated the noise robustness of the three algorithms described above by applying standard Gaussian noise at different Signal-to-Noise-Ratio (SNR) levels to the model responses during the test phase.

#### Model Generalization Evaluation

As discussed in (Gouwens et al., 2018), some combinations of predicted parameters can well perform under the stimulus protocol presented in the training phase, but function ill when applied to an unseen stimulus protocol. So, we also evaluated the generalization ability of the three algorithms to unseen stimulus protocols.

### Experimental Data Source and Pre-Training Configurations

All of the experimental recording data come from Allen Institute Cell Types Database(Allen Cell Types Database, 2016). We chose the experimental data from mouse spiny neurons. To perform the experiments in Figure 3, we first sampled 800K training data based on equation 7 and 8 using a 130pA long square stimulus protocol. During the stimulation, we only used the stimulus protocol from the 45000^th^ to the 115000^th^ time step and then downsampled it at the ratio of 10. We further did data washing to the original sampled training data by eliminating those whose firing frequency is higher than 36Hz or less than 5Hz, or the total number of spikes is less than 1 or more than 36, and which emit spontaneous spikes before the stimulus onset. The final size of the dataset for training and validation is 800K and the ratio of training and validation size is 99:1. The initial learning rate is set to be 10^-5^.

### SNPE Configurations

For both SNPE-alone and the combination of GNE and SNPE, we chose SNPEC to perform the inference, the density estimator is a 2-component mixture of Gaussian. There are 5 transforms in the SNPEC and the initial learning rate is set to be 10^-4^. There are 2 rounds of training in both cases. For SNPE-alone, the hand-crafted features are the same as those in (Gonçalves et al., 2020). Z-score transformation is applied to both the input summary features and the output parameters. As for the combination of GNE and SNPE, we used the pre-trained model on 800K synthesized training data and eliminated the FC layers. Z-score transformation is not applied to neither the raw response inputs nor the predicted parameters.

## Supplementary

### GNE Can Well Perform across a Wide Range of Neuronal Models

#### Izhikevich Model

Izhikevich neuronal model(Izhikevich, 2003) is a phenomenological model, which can describe a large category of neurons. This model has four parameters, namely *a*, *b*, *c*, *d*, whose lower bounds are [0, 0, 90, 0] and upper bounds are [0.1, 0.4, 40, 5].

In Figure S4a and b, we compared three metrics versus prediction accuracy. In terms of MAE of parameters, we achieved 3.7-6.4 times fewer errors than CNN. Under the training chirp stimulus protocol, the correlation between predicted model responses by GNE and the groundtruth is 2.2 times higher than that by CNN, and the MAE between the responses of GNE is 6.2 times less. As for the model generalization to unseen stimulus protocols, GNE outperformed CNN and MOO in all the stimulus protocols studies, and the results are presented in Figure S5.

In Figure S4c, we showed that GNE is more robust against Gaussian noise under nearly all levels of SNR on the Izhikevich neuronal model.

#### HH Ball-Stick Model

The HH ball-stick model we use contains a simplified soma and dendrite, with the same settings as in (Ben-Shalom et al., 2019). There are 8 parameters in this case, namely the maximum conductance of Na, K, Ca, L channels. In the training and test dataset, the maximum conductance of the L channel of the soma and the dendrite is always the same, but we do not manually control the output of GNE to make their values the same. The lower bounds of the parameters are [250, 5, 0.75, 0.0003, 250, 5, 0.75, 0.0003], the upper bounds are [1000, 20, 3, 0.0009, 1000, 20, 3, 0.0009].

As shown in Figure S6a, it can be seen that all the predictions of maximum conductance of K channel at dendrite except that by GNE are more accurate than the baseline, with 3.3 to 8.3 times less error. The correlations between the predicted responses and the groundtruth don’t show a significant difference between GNE and the baseline under both training chirp stimulus protocols and the unseen protocols. Under training chirp stimulus, both the correlation and MAE between predicted responses and the groundtruth are worse than those of the baseline, however, we achieved better performance under all of the unseen stimulus protocols (results shown in Figure S7).

In Figure S6c, we evaluated the noise robustness of GNE and the baseline. It can be clearly seen that GNE’s prediction of parameters gives higher accuracy, almost for all parameters and at all SNR levels

#### Mainen-Sejnowski Model: L4 Stellate Neuron and L5 Pyramidal Neuron

Mainen-Sejnowski model(Mainen & Sejnowski, 1996) is a biophysically detailed model with dendritic morphologies. There are 10 free parameters in this model that need estimating, namely the maximum conductance of Na^+^, K^+^, Ca^2+^ channels at soma, distal dendrite and basal dendrite, and the maximum conductance of leaky channel at soma. The lower boundsofthe parametersare [10,15000,10,1000,100,0.15, 0.05,1.5,0.3,1/60000], andtheupper bounds of the parameters are [250,80000,60000,5000,2000,0.6,0.2,6,1.5, 1/15000]. We chose two categories of neurons: L4 stellate neuron and L5 pyramidal neuron to perform the task.

Figure S8a and b and Figure S10a and b show the results of the L4 stellate neuron and the L5 pyramidal neuron. We first evaluated the prediction accuracy of predicted parameters and predicted model responses. GNE achieves 1.1-4.0 times less prediction MAE than CNN. As for the predicted model responses, both of the correlation and the MAE between the responses, GNE showed significant improvement over CNN.

Moreover, as shown in Figure S8c and Figure S10d, GNE is not sensitive to the Gaussian noise applied to the model responses for nearly all parameters except the prediction of Km^+^ channel due to its poor prediction accuracy.

---

**Figure S1:**
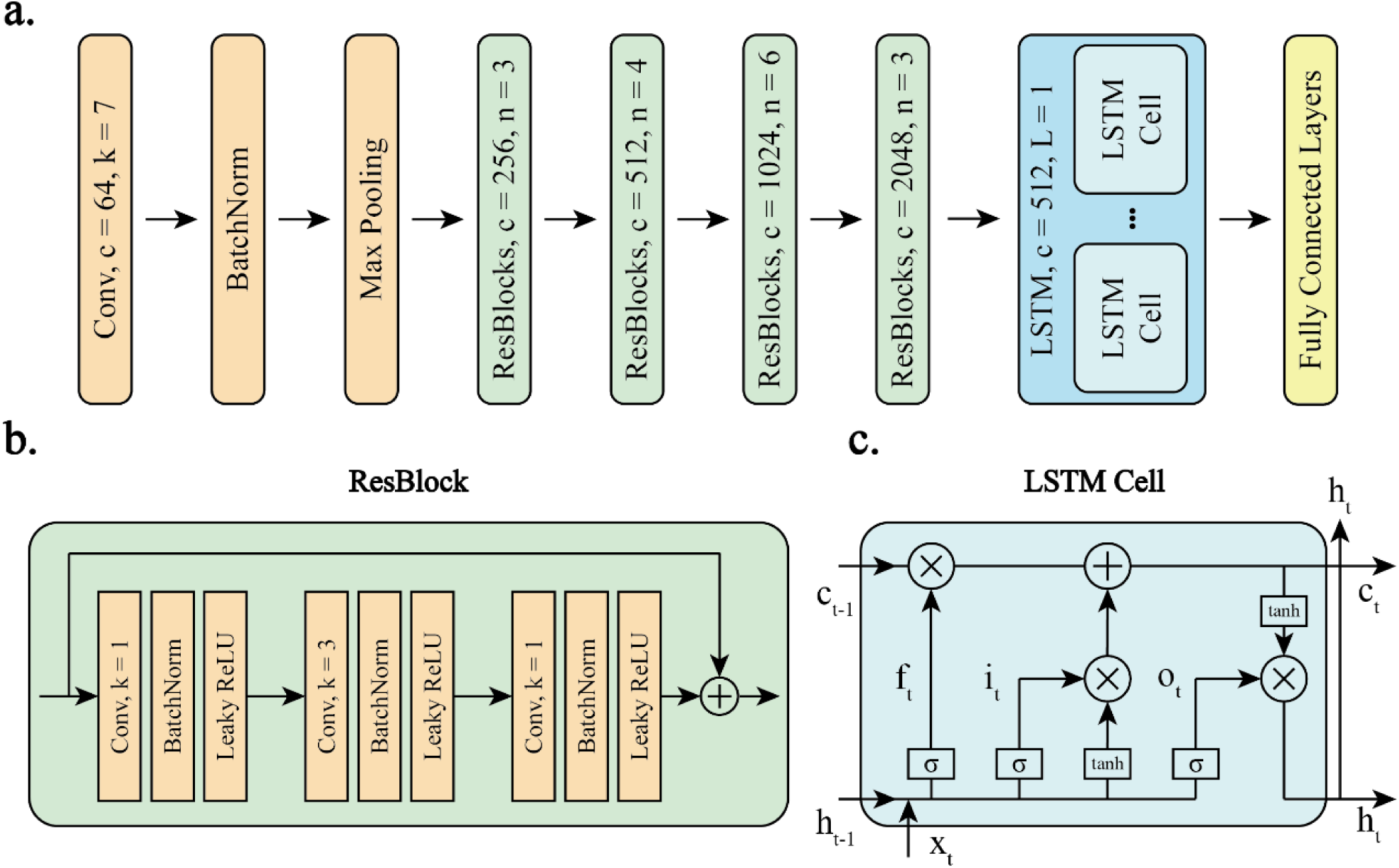
**a** Overall Deep Neural Network Architecture: each rectangular demonstrates a deep learning unit, where *c* means the dimension of output channels, *n* means the number of units in the rectangular, *L* means the number of layers. **b & c** Architecture and computation flow of residual block (ResBlock) and LSTM cell correspondingly, where each arrow means data transfer, *k* and BatchNorm in **b** mean the size of kernel of the convolution operator means the batch normalization layer, σ and *t* in c mean sigmoid function and the current time step.

**Figure S2:**
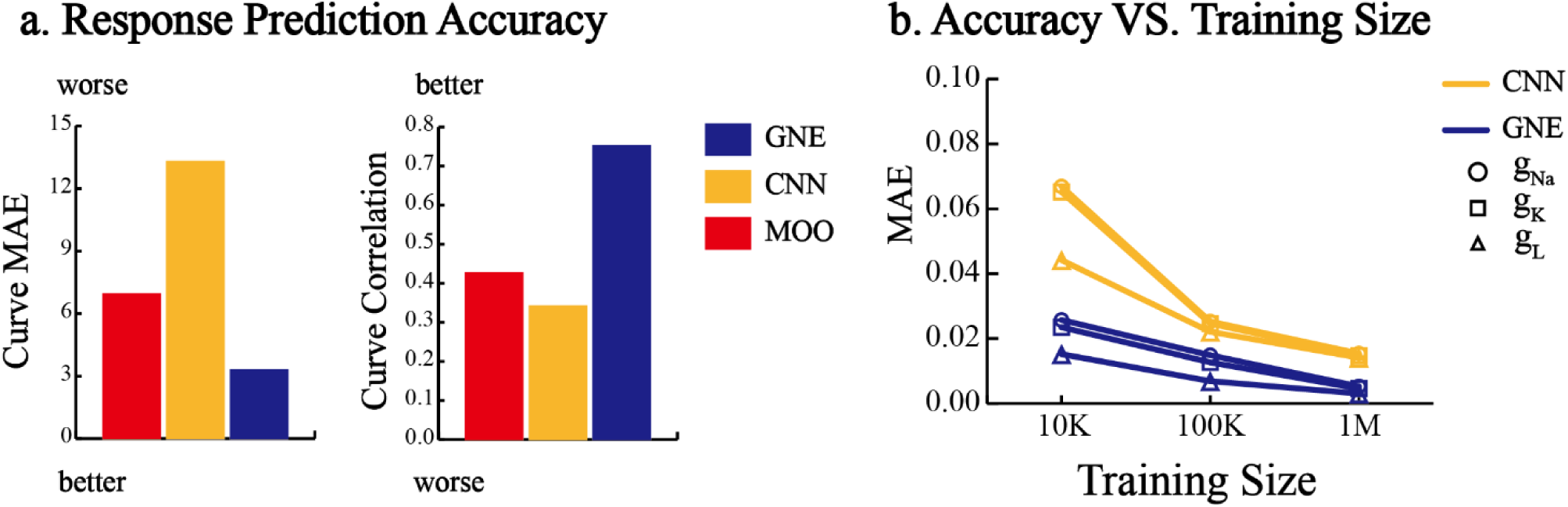
**a** Performance against MOO and CNN on Single HH Neuronal Model: accuracy of the predicted model responses under chirp stimulus protocol. **b** The relationship between prediction accuracy and training size for CNN and GNE.

**Figure S3:**
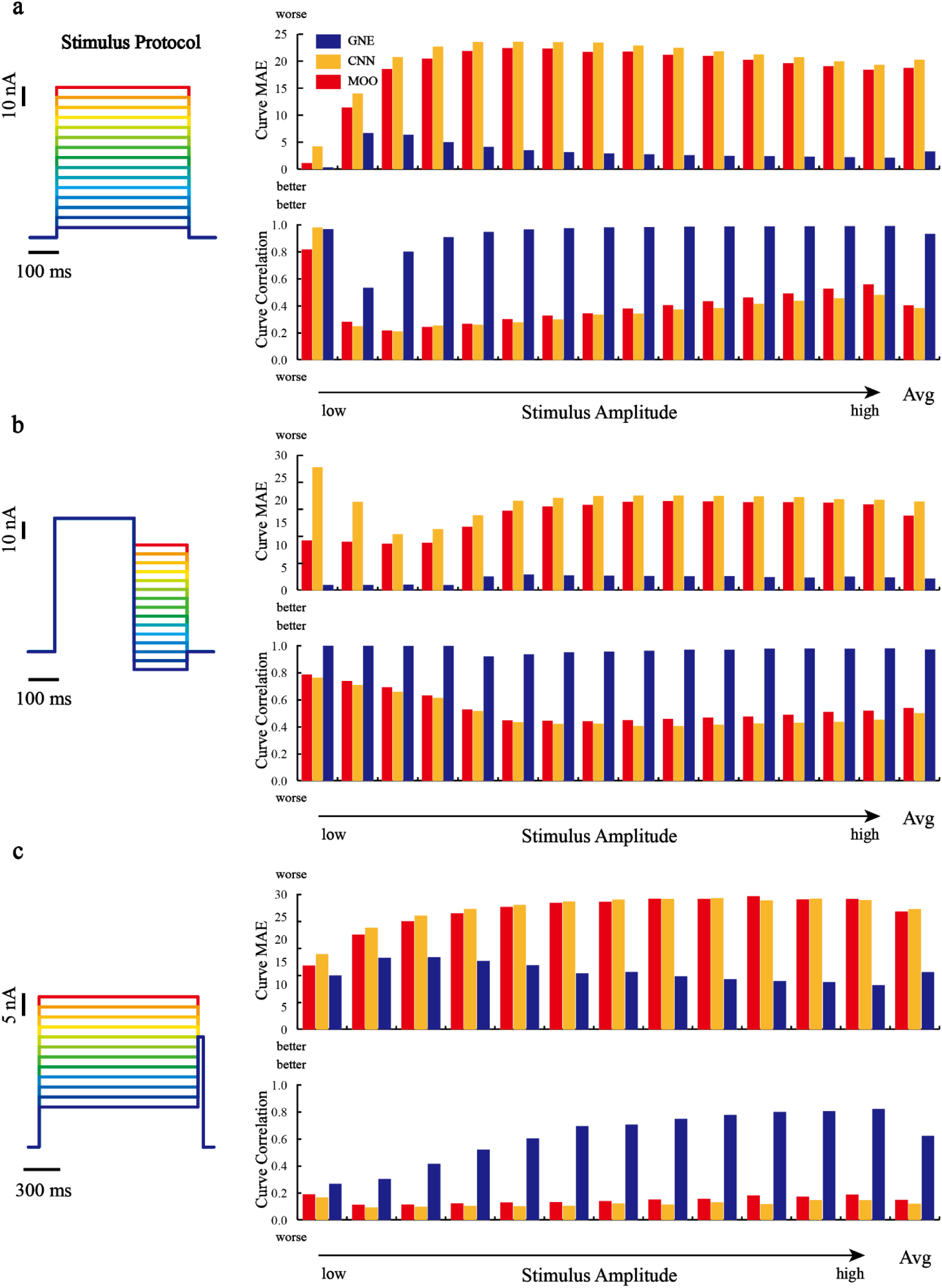
HH Neuronal Model Generalization Given Stimulus Protocols with Different Amplitude. Left: stimulus protocols. Right: MAE and Pearson correlation coefficient between predicted model responses and the groundtruth.

**Figure S4:**
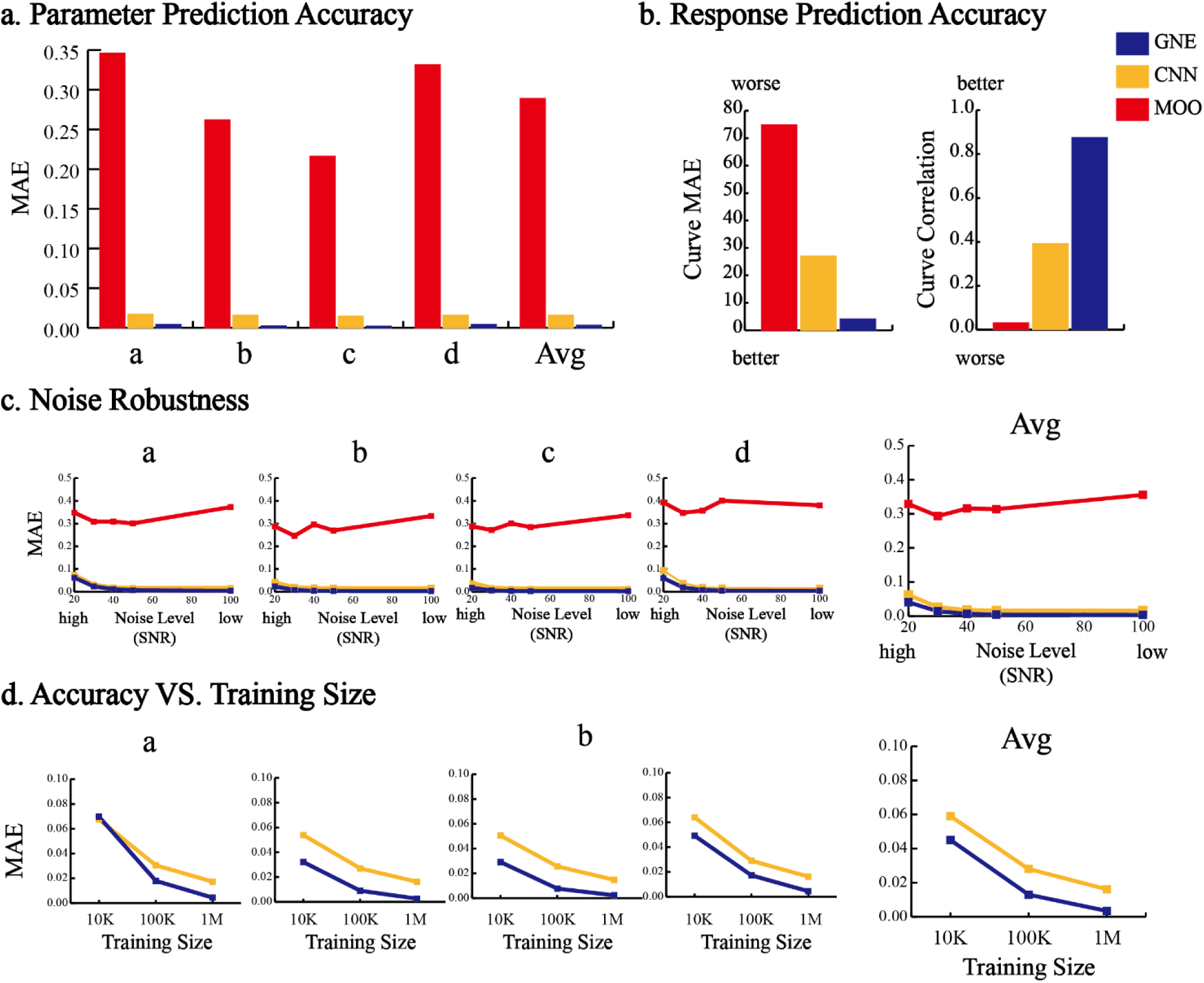
Performance against MOO and CNN on Izhikevich Neuronal Model. **a** Accuracy of the predicted parameters. **b** Accuracy of the predicted model responses under chirp stimulus protocol. **c** Noise robustness test compared with the baseline.

**Figure S5:**
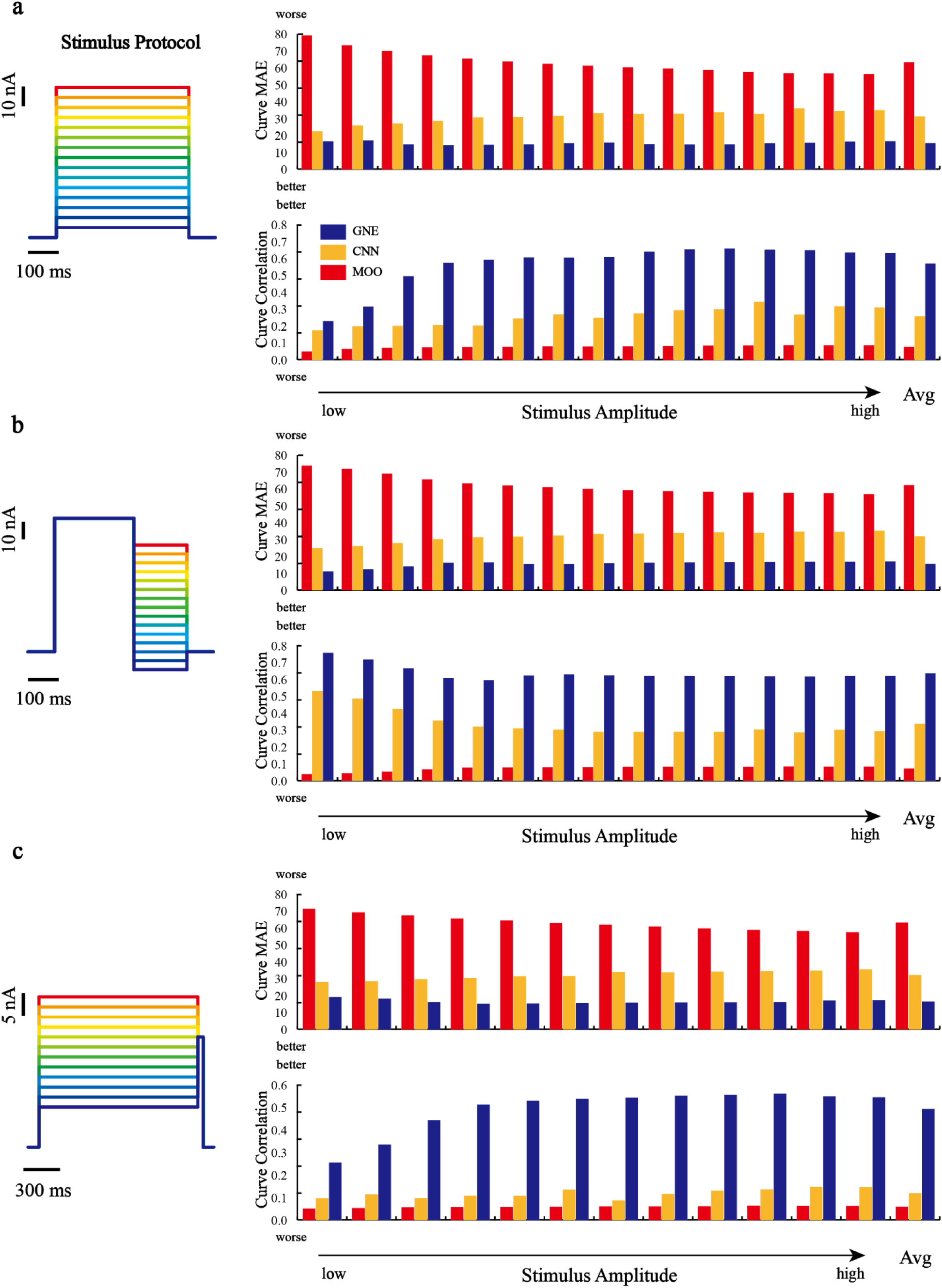
Izhikevich Neuronal Model Generalization Given Stimulus Protocols with Different Amplitude. Left: stimulus protocols. Right: MAE and Pearson correlation coefficient between predicted model responses and the groundtruth.

**Figure S6:**
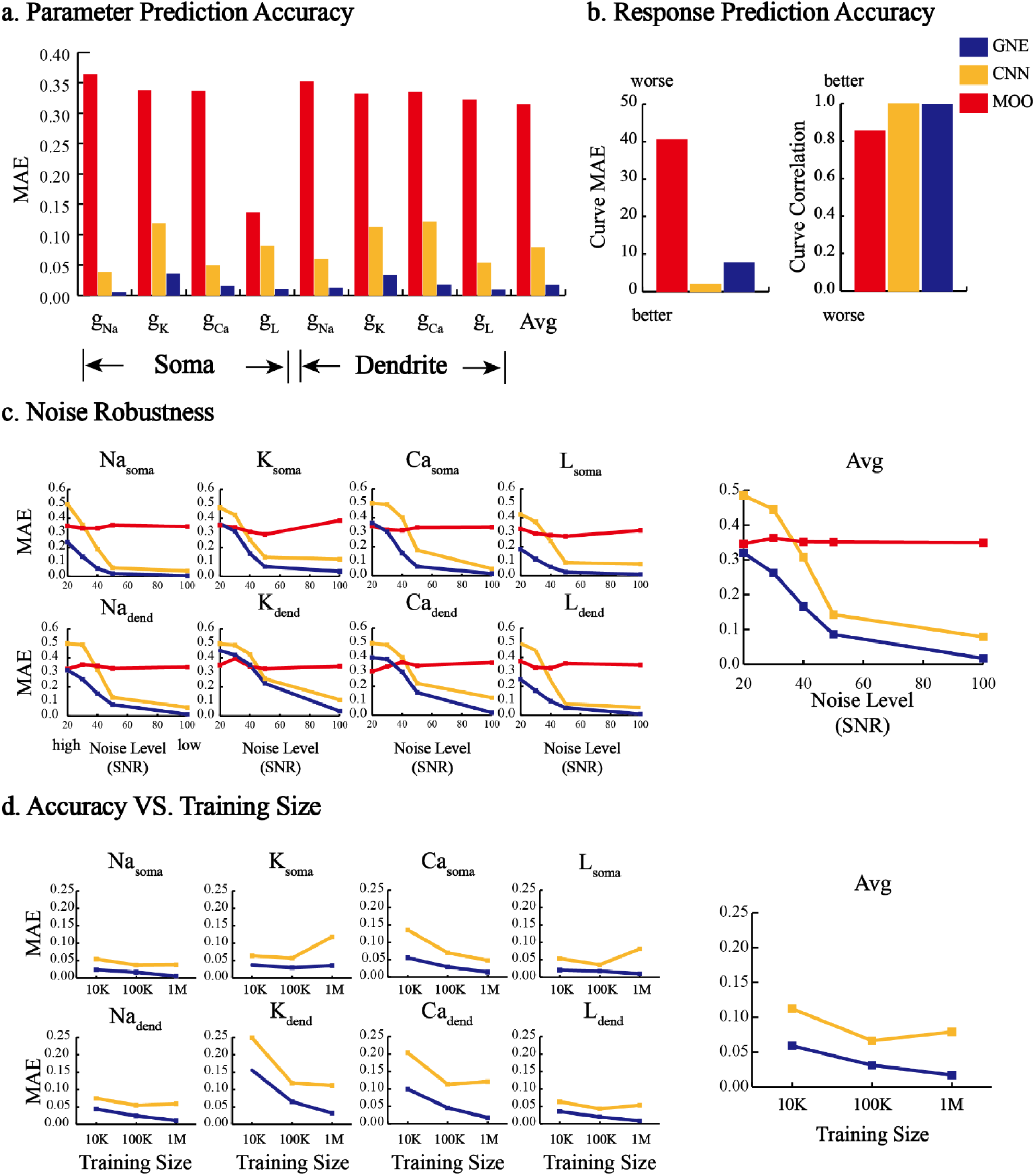
Performance against MOO and CNN on HH Ball-Stick Neuronal Model. **a** Accuracy of the predicted parameters. **b** Accuracy of the predicted model responses under chirp stimulus protocol. **c** Noise robustness test compared with the baseline. **d** The relationship between the accuracy of parameter prediction and the training size between GNE and CNN.

**Figure S7:**
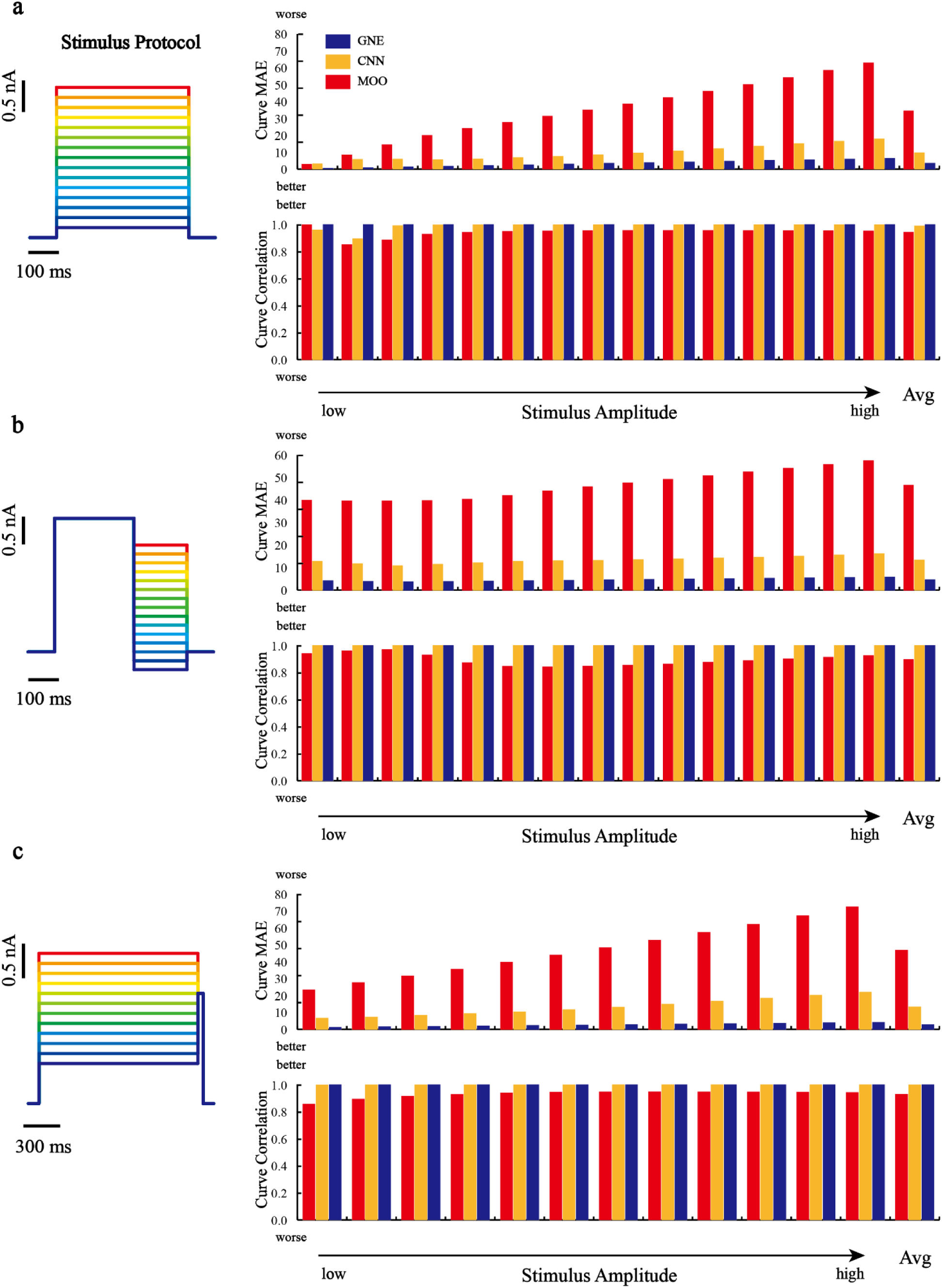
HH Ball-Stick Neuronal Model Generalization Given Stimulus Protocols with Different Amplitude. Left: stimulus protocols. Right: MAE and Pearson correlation coefficient between predicted model responses and the groundtruth.

**Figure S8:**
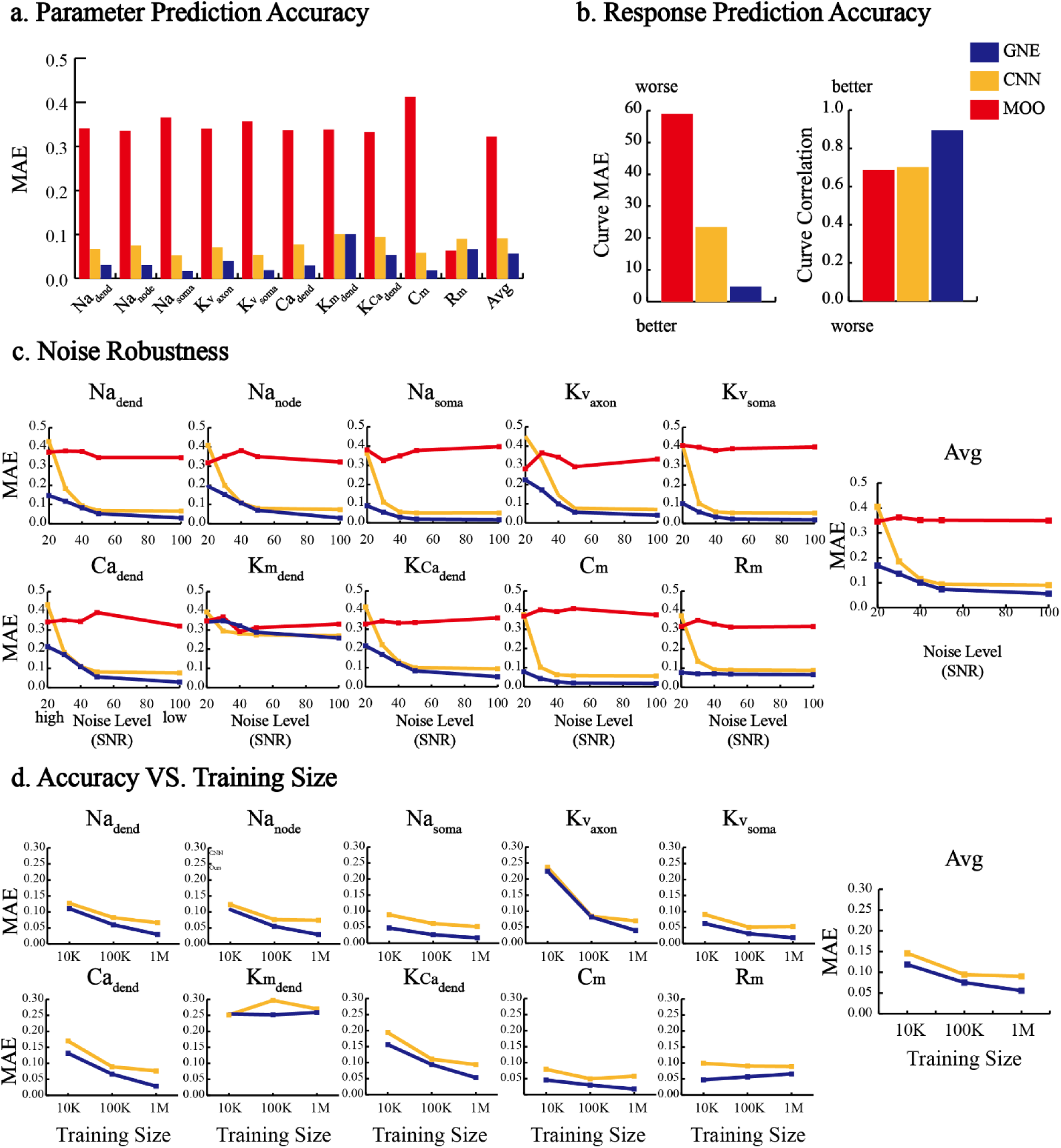
Performance against MOO and CNN on L4 Stellate Neuronal Model. **a** Accuracy of the predicted parameters. **b** Accuracy of the predicted model responses under chirp stimulus protocol. **c** Noise robustness test compared with the baseline. **d** The relationship between the accuracy of parameter prediction and the training size between GNE and CNN.

**Figure S9:**
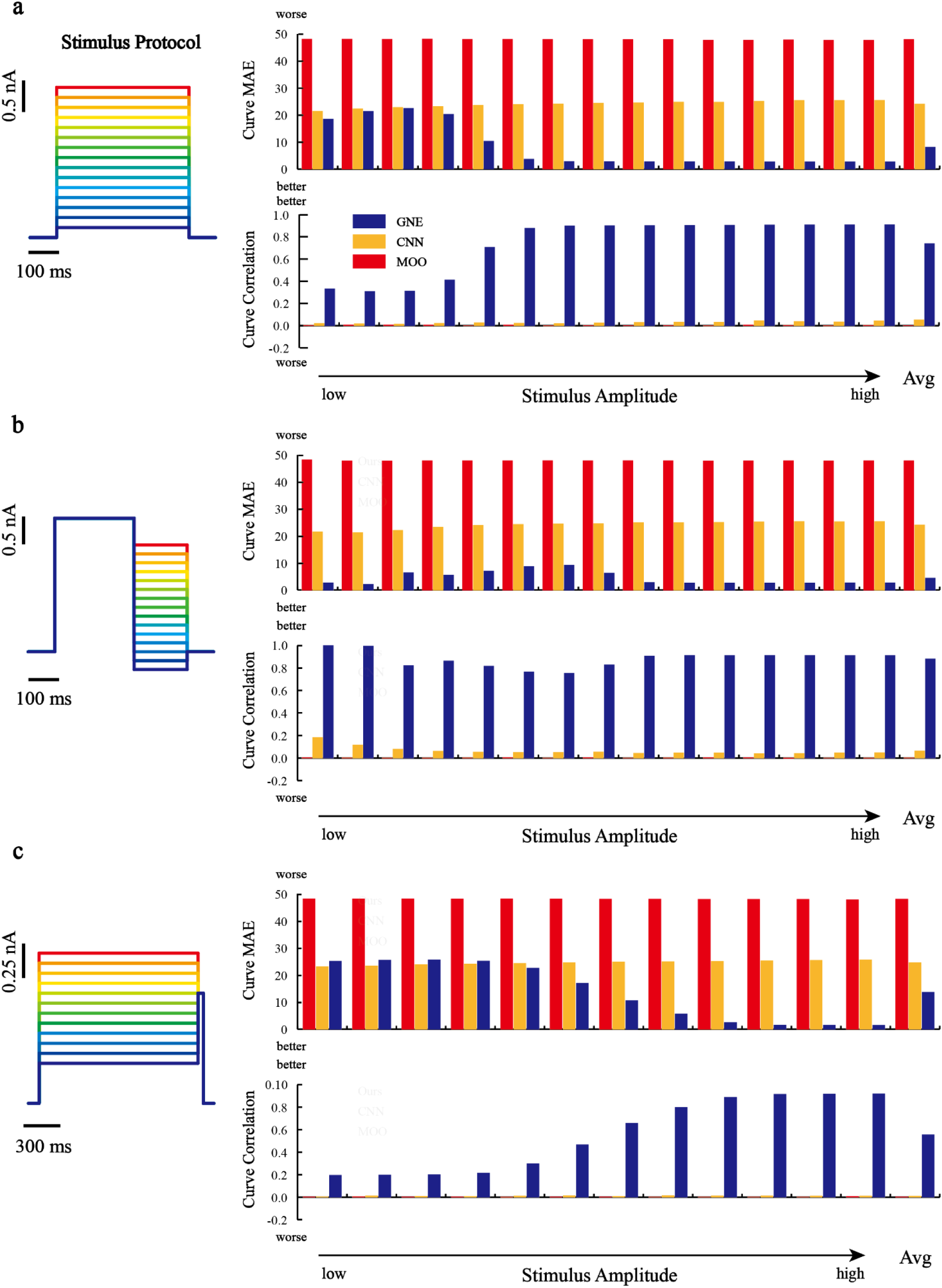
L4 Stellate Neuronal Model Generalization Given Stimulus Protocols with Different Amplitude. Left: stimulus protocols. Right: MAE and Pearson correlation coefficient between predicted model responses and the groundtruth.

**Figure S10:**
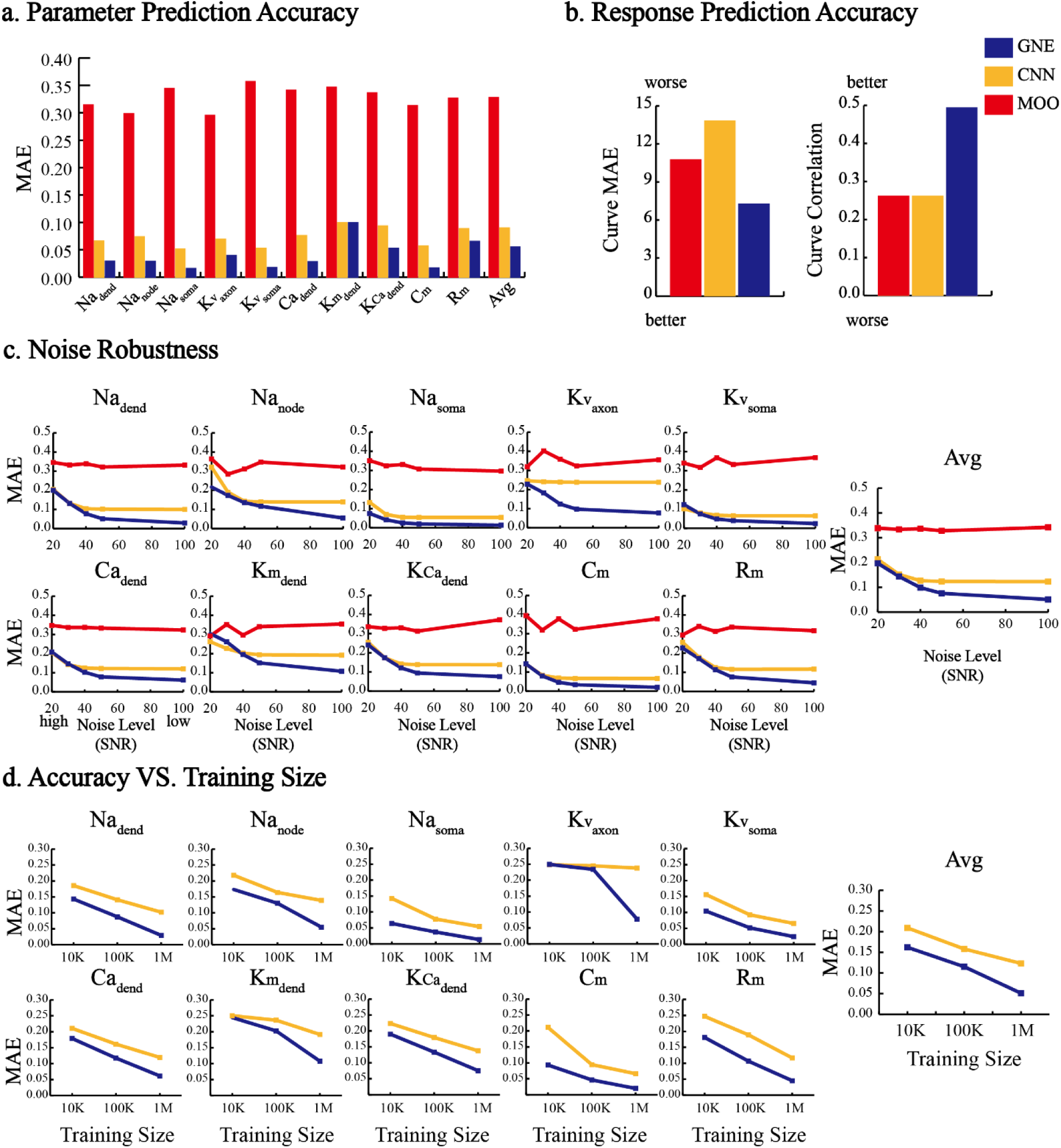
Performance against MOO and CNN on L5 Pyramidal Neuronal Model. **a** Accuracy of the predicted parameters. **b** Accuracy of the predicted model responses under chirp stimulus protocol. **c** Noise robustness test compared with the baseline. **d** The relationship between the accuracy of parameter prediction and the training size between GNE and CNN.

**Figure S11:**
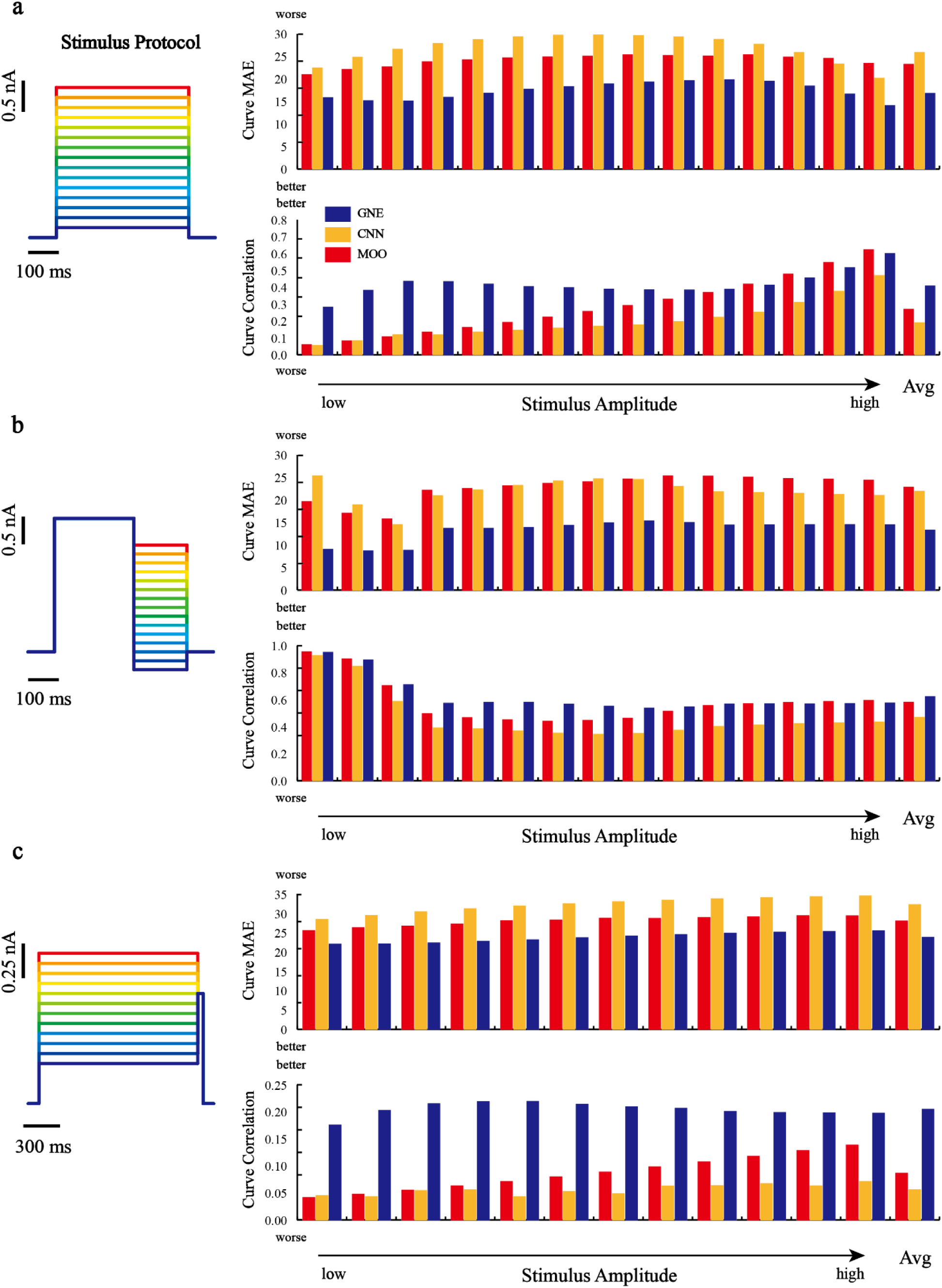
L5 Pyramidal Model Generalization Given Stimulus Protocols with Different Amplitude. Left: stimulus protocols. Right: MAE and Pearson correlation coefficient between predicted model responses and the groundtruth.

**Figure S12:**
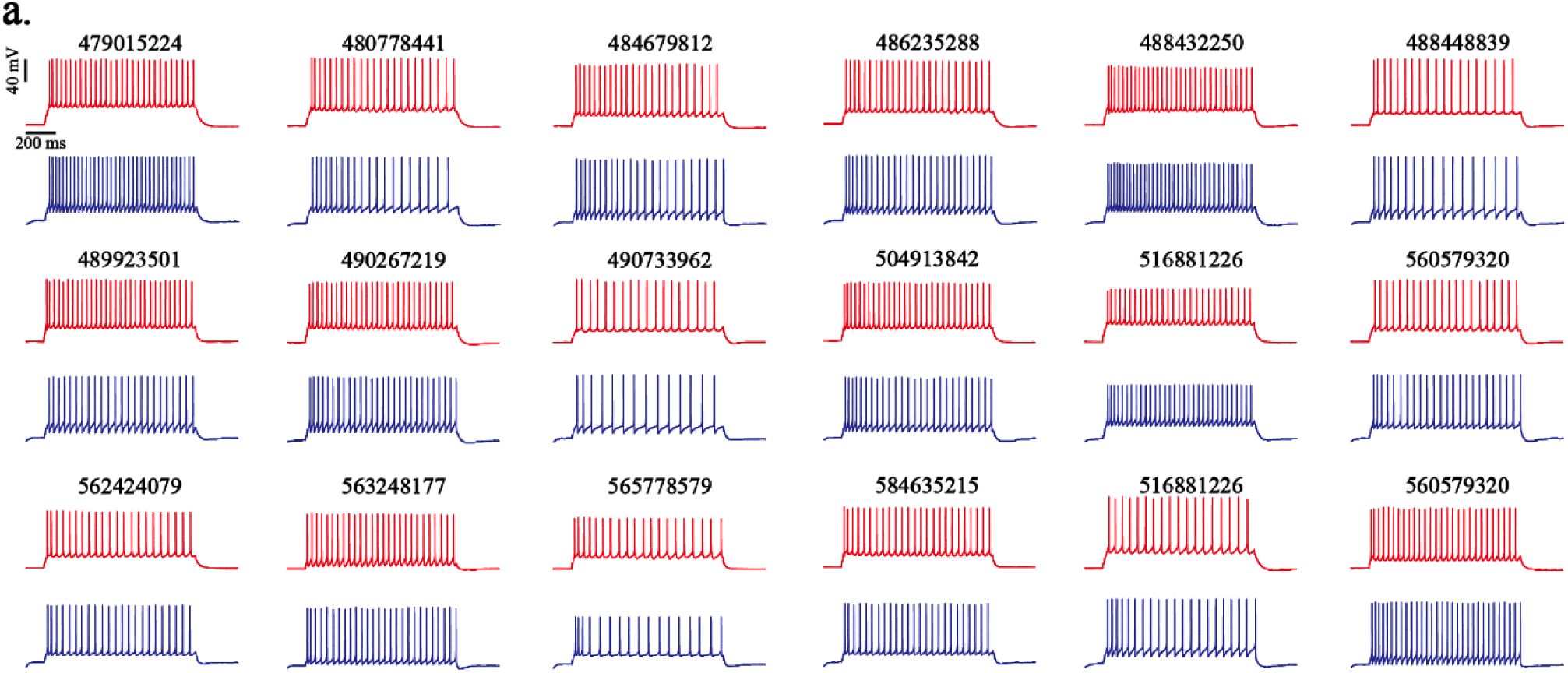
Results for direct estimation on experimental data from Allen Institute(Allen Cell Types Database, 2016). The numbers denote the specimen ID, and all experimental neural responses are obtained from step current with amplitude 130 pA.

**Figure S13:**
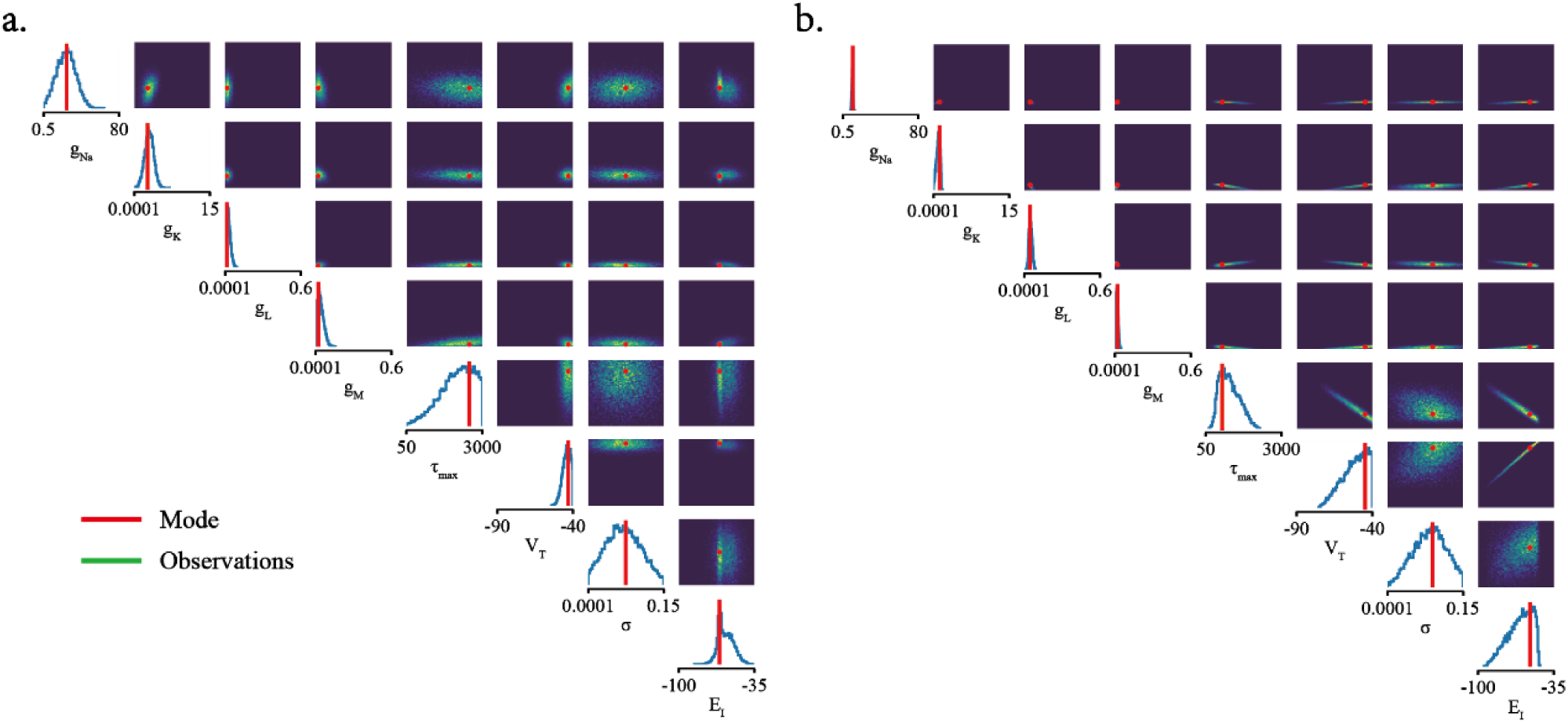
**a** The complete predicted marginal distribution density map shown in Figure 3c, where the green dots are the samples from the posterior distribution and the red dot represents the mode position. **b** The complete predicted marginal distribution density map shown in Figure 3d.

**Figure S14:**
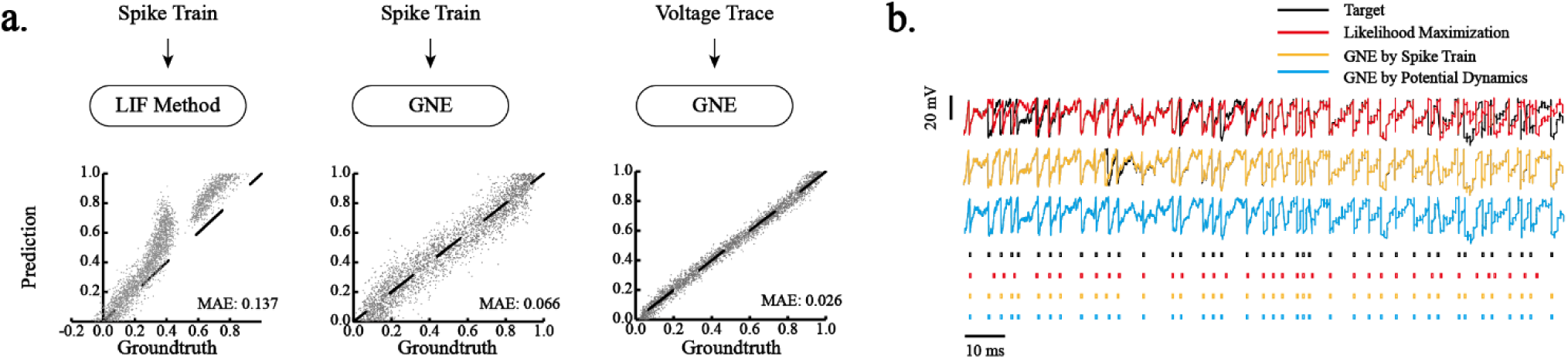
**a** Performance comparison with the I&F method(Ladenbauer et al., 2019). **b** An example of the predicted circuitry dynamics and spike trains of a selected neuron by the three methods in **a**.

**Figure S15:**
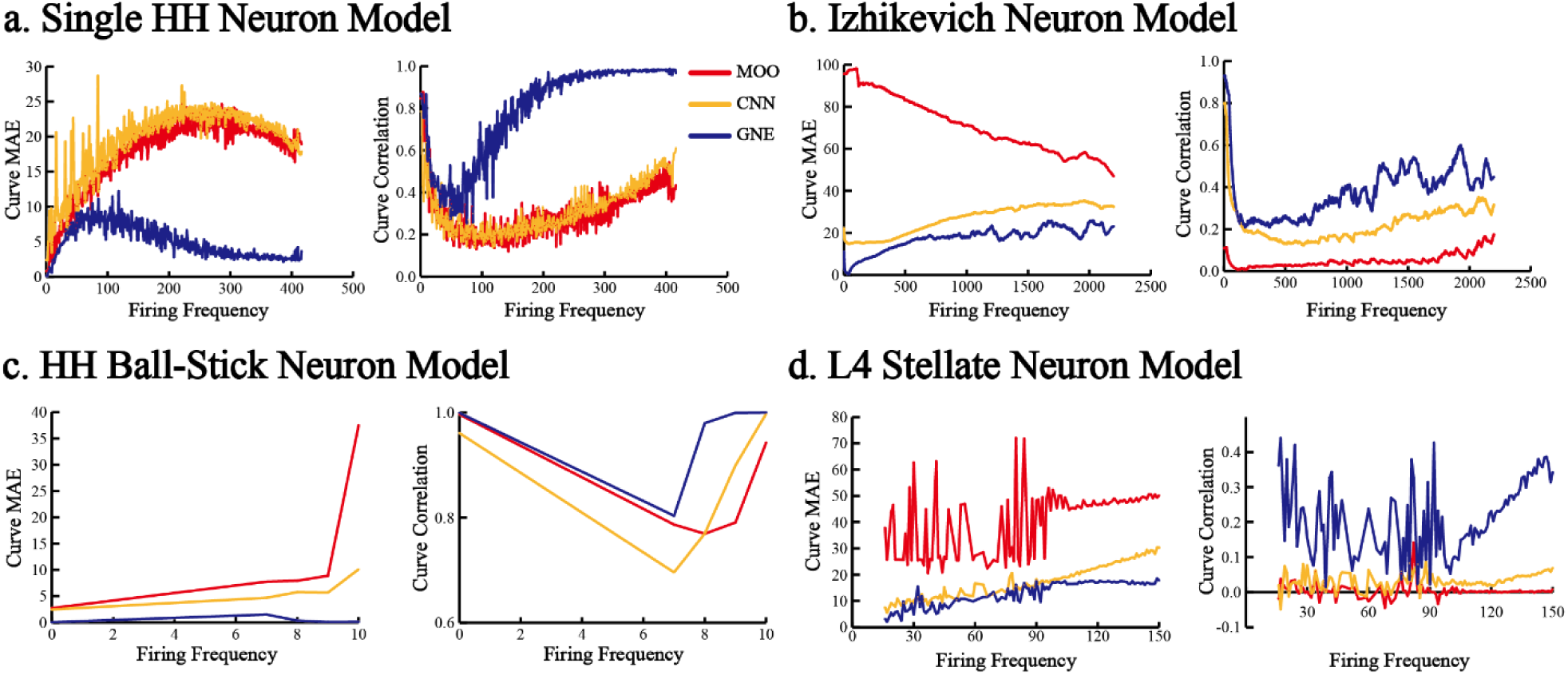
Scores of Two Response Prediction Criteria vs. Neuronal Model Firing Frequency.

## References

Abbott, L. F. (1999). Lapicque’s introduction of the integrate-and-fire model neuron (1907). Brain research bulletin, 50(5-6), 303–304 %@ 0361-9230. doi:10.1016/S0361-9230(99)00161-6

Abbott, L. F. (2008). Theoretical neuroscience rising. Neuron, 60(3), 489–495 %@ 0896-6273. doi:10.1016/j.neuron.2008.10.019

Adesnik, H., Bruns, W., Taniguchi, H., Huang, Z. J., & Scanziani, M. (2012). A neural circuit for spatial summation in visual cortex. Nature, 490(7419), 226–231. doi:10.1038/nature11526

Allen Cell Types Database. (2016). Retrieved from: http://celltypes.brain-map.org

Atiya, N. A., Rañó, I., Prasad, G., & Wong-Lin, K. (2019). A neural circuit model of decision uncertainty and change-of-mind. Nature communications, 10(1), 1–12. doi:10.1038/s41467-019-10316-8

Baden, T., Esposti, F., Nikolaev, A., & Lagnado, L. (2011). Spikes in retinal bipolar cells phase-lock to visual stimuli with millisecond precision. Current Biology, 21(22), 1859–1869 %@ 0960-9822. doi:10.1016/j.cub.2011.09.042

Bahl, A., Stemmler, M. B., Herz, A. V., & Roth, A. (2012). Automated optimization of a reduced layer 5 pyramidal cell model based on experimental data. Journal of neuroscience methods, 210(1), 22–34. doi:10.1016/j.jneumeth.2012.04.006

Ben-Shalom, R., Balewski, J., Siththaranjan, A., Baratham, V., Kyoung, H., Kim, K. G., … Bouchard, K. E. (2019). Inferring neuronal ionic conductances from membrane potentials using CNNs. bioRxiv, 727974.

Billeh, Y. N., Cai, B., Gratiy, S. L., Dai, K., Iyer, R., Gouwens, N. W., … Olsen, S. R. (2020). Systematic integration of structural and functional data into multi-scale models of mouse primary visual cortex. Neuron, 106(3), 388–403.e318. doi:10.1016/j.neuron.2020.01.040

Briggman, K. L., Helmstaedter, M., & Denk, W. (2011). Wiring specificity in the direction-selectivity circuit of the retina. Nature, 471(7337), 183–188. doi:10.1038/nature09818

Brown, T. B., Mann, B., Ryder, N., Subbiah, M., Kaplan, J., Dhariwal, P., … Askell, A. (2020). Language models are few-shot learners. arXiv preprint arXiv:2005.14165.

Bush, K., Knight, J., & Anderson, C. (2005). Optimizing conductance parameters of cortical neural models via electrotonic partitions. Neural networks, 18(5-6), 488–496. doi:10.1016/j.neunet.2005.06.038

Cadwell, C. R., Palasantza, A., Jiang, X., Berens, P., Deng, Q., Yilmaz, M., … Tolias, K. F. (2016). Electrophysiological, transcriptomic and morphologic profiling of single neurons using Patch-seq. Nature biotechnology, 34(2), 199–203. doi:10.1038/nbt.3445

Carnevale, N. T., & Hines, M. L. (2006). The NEURON book: Cambridge University Press.

Clay, J. R., & Shrier, A. (1999). On the role of subthreshold dynamics in neuronal signaling. Journal of theoretical biology, 197(2), 207–216 %@ 0022-5193. doi:10.1006/jtbi.1998.0867

David, G., Marcel, N., & Jakob, M. (2019). Automatic Posterior Transformation for Likelihood-Free Inference. In C. Kamalika & S. Ruslan (Eds.), Proceedings of the 36th International Conference on Machine Learning (Vol. 97, pp. 2404--2414). Proceedings of Machine Learning Research: PMLR.

Destexhe, A., Mainen, Z. F., & Sejnowski, T. J. (1998). Kinetic models of synaptic transmission. Methods in neuronal modeling, 2, 1–25.

Druckmann, S., Banitt, Y., Gidon, A. A., Schürmann, F., Markram, H., & Segev, I. (2007). A novel multiple objective optimization framework for constraining conductance-based neuron models by experimental data. Frontiers in neuroscience, 1, 1. doi:10.3389/neuro.01.1.1.001.2007

Druckmann, S., Berger, T. K., Hill, S., Schürmann, F., Markram, H., & Segev, I. (2008). Evaluating automated parameter constraining procedures of neuron models by experimental and surrogate data. Biological cybernetics, 99(4-5), 371. doi:10.1007/s00422-008-0269-2

Du, K., Wu, Y.-W., Lindroos, R., Liu, Y., Rózsa, B., Katona, G., … Kotaleski, J. H. (2017). Cell-type– specific inhibition of the dendritic plateau potential in striatal spiny projection neurons. Proceedings of the National Academy of Sciences, 114(36), E7612–E7621 %@ 0027-8424. doi:10.1073/pnas.1704893114

Ebner, C., Clopath, C., Jedlicka, P., & Cuntz, H. (2019). Unifying Long-Term Plasticity Rules for Excitatory Synapses by Modeling Dendrites of Cortical Pyramidal Neurons. Cell reports, 29(13), 4295–4307. e4296 %@ 2211-1247. doi:10.1016/j.celrep.2019.11.068

Egger, S. W., Le, N. M., & Jazayeri, M. (2020). A neural circuit model for human sensorimotor timing. Nature communications, 11(1), 1–14. doi:10.1038/s41467-020-16999-8

Eliasmith, C., Gosmann, J., & Choo, X. (2016). BioSpaun: A large-scale behaving brain model with complex neurons. arXiv preprint arXiv:1602.05220. doi:10.1371/journal.pcbi.0020094

Endo, D., Kobayashi, R., Bartolo, R., Averbeck, B. B., Sugase-Miyamoto, Y., Hayashi, K., … Shinomoto, S. (2020). CoNNECT: Convolutional Neural Network for Estimating synaptic Connectivity from spike Trains. bioRxiv. doi:10.1101/2020.05.05.078089

Franconville, R., Beron, C., & Jayaraman, V. (2018). Building a functional connectome of the Drosophila central complex. Elife, 7, e37017. doi:10.7554/eLife.37017.001

Gamboa, J. C. B. (2017). Deep learning for time-series analysis. arXiv preprint arXiv:1701.01887.

Gerken, W. C., Purvis, L., & Butera, R. J. (2006). Genetic algorithm for optimization and specification of a neuron model. Neurocomputing, 69(10-12), 1039–1042. doi:10.1016/j.neucom.2005.12.041

Gerstner, W., & Naud, R. (2009). How good are neuron models? Science, 326(5951), 379–380. doi:10.1126/science.1181936

Goaillard, J.-M., Taylor, A. L., Schulz, D. J., & Marder, E. (2009). Functional consequences of animal-to-animal variation in circuit parameters. Nature neuroscience, 12(11), 1424–1430. doi:10.1038/nn.2404

Gonçalves, P. J., Lueckmann, J.-M., Deistler, M., Nonnenmacher, M., Öcal, K., Bassetto, G., … Macke, J. H. (2020). Training deep neural density estimators to identify mechanistic models of neural dynamics. Elife, 9, e56261. doi:10.7554/eLife.56261

Gouwens, N. W., Berg, J., Feng, D., Sorensen, S. A., Zeng, H., Hawrylycz, M. J., … Arkhipov, A. (2018). Systematic generation of biophysically detailed models for diverse cortical neuron types. Nature communications, 9(1), 1–13. doi:10.1038/s41467-017-02718-3

Grienberger, C., & Konnerth, A. (2012). Imaging calcium in neurons. Neuron, 73(5), 862–885. doi:10.1016/j.neuron.2012.02.011

Gurkiewicz, M., & Korngreen, A. (2007). A numerical approach to ion channel modelling using whole-cell voltage-clamp recordings and a genetic algorithm. PLoS Comput Biol, 3(8), e169. doi:10.1371/journal.pcbi.0030169

He, K., Zhang, X., Ren, S., & Sun, J. (2016). Deep residual learning for image recognition. Paper presented at the Proceedings of the IEEE conference on computer vision and pattern recognition.

Hendrickson, E. B., Edgerton, J. R., & Jaeger, D. (2011). The use of automated parameter searches to improve ion channel kinetics for neural modeling. Journal of computational neuroscience, 31(2), 329–346. doi:10.1007/s10827-010-0312-x

Hillenbrand, U. (2002). Subthreshold dynamics of the neural membrane potential driven by stochastic synaptic input. Physical Review E, 66(2), 021909. doi:10.1103/PhysRevE.66.021909

Hjorth, J. J. J., Kozlov, A., Carannante, I., Nylén, J. F., Lindroos, R., Johansson, Y., … Silberberg, G. (2020). The microcircuits of striatum in silico. Proceedings of the National Academy of Sciences, 117(17), 9554–9565 %@ 0027-8424. doi:10.1073/pnas.2000671117

Hochreiter, S., & Schmidhuber, J. (1997). Long short-term memory. Neural computation, 9(8), 1735–1780. doi:10.1162/neco.1997.9.8.1735

Hodgkin, A. L., & Huxley, A. F. (1952). A quantitative description of membrane current and its application to conduction and excitation in nerve. The Journal of physiology, 117(4), 500. doi:10.1113/jphysiol.1952.sp004764

Huys, Q. J., Ahrens, M. B., & Paninski, L. (2006). Efficient estimation of detailed single-neuron models. Journal of neurophysiology, 96(2), 872–890. doi:10.1152/jn.00079.2006

Huys, Q. J., & Paninski, L. (2009). Smoothing of, and parameter estimation from, noisy biophysical recordings. PLoS Comput Biol, 5(5), e1000379. doi:10.1371/journal.pcbi.1000379

Huys, Q. J. M., Maia, T. V., & Frank, M. J. (2016). Computational psychiatry as a bridge from neuroscience to clinical applications. Nature neuroscience, 19(3), 404 %@ 1546-1726. doi:10.1038/nn.4238

Izhikevich, E. M. (2003). Simple model of spiking neurons. IEEE Transactions on neural networks, 14(6), 1569–1572. doi:10.1109/TNN.2003.820440

Jolivet, R., Kobayashi, R., Rauch, A., Naud, R., Shinomoto, S., & Gerstner, W. (2008). A benchmark test for a quantitative assessment of simple neuron models. Journal of neuroscience methods, 169(2), 417–424. doi:10.1016/j.jneumeth.2007.11.006

Jolivet, R., Roth, A., Schürmann, F., Gerstner, W., & Senn, W. (2008). Special issue on quantitative neuron modeling. Biological cybernetics, 99(4-5), 237–239.

Keren, N., Peled, N., & Korngreen, A. (2005). Constraining compartmental models using multiple voltage recordings and genetic algorithms. Journal of neurophysiology. doi:10.1152/jn.00408.2005

Kingma, D. P., & Ba, J. (2014). Adam: A method for stochastic optimization. arXiv preprint arXiv:1412.6980.

Kobayashi, R., Kurita, S., Kurth, A., Kitano, K., Mizuseki, K., Diesmann, M., … Shinomoto, S. (2019). Reconstructing neuronal circuitry from parallel spike trains. Nature communications, 10(1), 1–13. doi:10.1038/s41467-019-12225-2

Kobayashi, R., & Shinomoto, S. (2007). Predicting spike times from subthreshold dynamics of a neuron.

Koch, C., & Segev, I. (1998). Methods in neuronal modeling: from ions to networks: MIT press.

Krizhevsky, A., Sutskever, I., & Hinton, G. E. (2012). Imagenet classification with deep convolutional neural networks. Paper presented at the Advances in neural information processing systems.

Ladenbauer, J., McKenzie, S., English, D. F., Hagens, O., & Ostojic, S. (2019). Inferring and validating mechanistic models of neural microcircuits based on spike-train data. Nature communications, 10(1), 1–17 %@ 2041-1723. doi:10.1038/s41467-019-12572-0

Larkum, M. E., Zhu, J. J., & Sakmann, B. (1999). A new cellular mechanism for coupling inputs arriving at different cortical layers. Nature, 398(6725), 338–341 %@ 1476-4687. doi:10.1038/18686

Latorre, R., Torres, J. J., & Varona, P. (2016). Interplay between subthreshold oscillations and depressing synapses in single neurons. PloS one, 11(1), e0145830 %@ 0141932-0146203. doi:10.1371/journal.pone.0145830

Lea, C., Vidal, R., Reiter, A., & Hager, G. D. (2016). Temporal Convolutional Networks: A Unified Approach to Action Segmentation, Cham %@ 978-3-319-49409-8.

Leshno, M., Lin, V. Y., Pinkus, A., & Schocken, S. (1993). Multilayer feedforward networks with a nonpolynomial activation function can approximate any function. Neural networks, 6(6), 861–867 %@ 0893-6080. doi:10.1016/S0893-6080(05)80131-5

Lueckmann, J.-M., Goncalves, P. J., Bassetto, G., Öcal, K., Nonnenmacher, M., & Macke, J. H. (2017). Flexible statistical inference for mechanistic models of neural dynamics.

Mainen, Z. F., & Sejnowski, T. J. (1996). Influence of dendritic structure on firing pattern in model neocortical neurons. Nature, 382(6589), 363–366. doi:10.1038/382363a0

Mann, K., Gallen, C. L., & Clandinin, T. R. (2017). Whole-brain calcium imaging reveals an intrinsic functional network in Drosophila. Current Biology, 27(15), 2389–2396. e2384. doi:10.1016/j.cub.2017.06.076

Markram, H., Muller, E., Ramaswamy, S., Reimann, M. W., Abdellah, M., Sanchez, C. A., … Arsever, S. (2015). Reconstruction and simulation of neocortical microcircuitry. Cell, 163(2), 456–492. doi:10.1016/j.cell.2015.09.029

Masoli, S., Rizza, M. F., Sgritta, M., Van Geit, W., Schürmann, F., & D’Angelo, E. (2017). Single neuron optimization as a basis for accurate biophysical modeling: the case of cerebellar granule cells. Frontiers in cellular neuroscience, 11, 71 %@ 1662-5102. doi:10.3389/fncel.2017.00071

McDougal, R. A., Morse, T. M., Carnevale, T., Marenco, L., Wang, R., Migliore, M., … Hines, M. L. (2017). Twenty years of ModelDB and beyond: building essential modeling tools for the future of neuroscience. Journal of computational neuroscience, 42(1), 1–10. doi:10.1007/s10827-016-0623-7

Meliza, C. D., Kostuk, M., Huang, H., Nogaret, A., Margoliash, D., & Abarbanel, H. D. (2014). Estimating parameters and predicting membrane voltages with conductance-based neuron models. Biological cybernetics, 108(4), 495–516. doi:10.1007/s00422-014-0615-5

Menon, V., Spruston, N., & Kath, W. L. (2009). A state-mutating genetic algorithm to design ion-channel models. Proceedings of the National Academy of Sciences, 106(39), 16829–16834. doi:10.1073/pnas.0903766106

Ness, T. V., Remme, M. W. H., & Einevoll, G. T. (2016). Active subthreshold dendritic conductances shape the local field potential. The Journal of physiology, 594(13), 3809–3825 %@ 0022-3751. doi:10.1113/JP272022

Neymotin, S. A., Suter, B. A., Dura-Bernal, S., Shepherd, G. M., Migliore, M., & Lytton, W. W. (2017). Optimizing computer models of corticospinal neurons to replicate in vitro dynamics. Journal of neurophysiology, 117(1), 148–162. doi:10.1152/jn.00570.2016

Oesterle, J., Behrens, C., Schroeder, C., Herrmann, T., Euler, T., Franke, K., … Berens, P. (2020). Bayesian inference for biophysical neuron models enables stimulus optimization for retinal neuroprosthetics. Elife, 9, e54997. doi:10.7554/eLife.54997

Otomo, K., Perkins, J., Kulkarni, A., Stojanovic, S., Roeper, J., & Paladini, C. A. (2020). Subthreshold repertoire and threshold dynamics of midbrain dopamine neuron firing in vivo. bioRxiv. doi:10.1101/2020.04.06.028829

Papamakarios, G., & Murray, I. (2016). Fast ε-free inference of simulation models with bayesian conditional density estimation. Paper presented at the Advances in neural information processing systems.

Papamakarios, G., Pavlakou, T., & Murray, I. (2017). Masked autoregressive flow for density estimation.

Petoe, M. A., Bradley, A. P., & Wilson, W. J. (2010). On chirp stimuli and neural synchrony in the suprathreshold auditory brainstem response. The Journal of the Acoustical Society of America, 128(1), 235–246 %@ 0001-4966. doi:10.1121/1.3436527

Pillow, J. W., Shlens, J., Paninski, L., Sher, A., Litke, A. M., Chichilnisky, E. J., & Simoncelli, E. P. (2008). Spatio-temporal correlations and visual signalling in a complete neuronal population. Nature, 454(7207), 995–999 %@ 1476-4687. doi:10.1038/nature07140

Podlaski, W. F., Seeholzer, A., Groschner, L. N., Miesenböck, G., Ranjan, R., & Vogels, T. P. (2017). Mapping the function of neuronal ion channels in model and experiment. Elife, 6, e22152. doi:10.7554/eLife.22152

Pozzorini, C., Mensi, S., Hagens, O., Naud, R., Koch, C., & Gerstner, W. (2015). Automated high-throughput characterization of single neurons by means of simplified spiking models. PLoS Comput Biol, 11(6), e1004275. doi:10.1371/journal.pcbi.1004275

Prinz, A. A., Billimoria, C. P., & Marder, E. (2003). Alternative to hand-tuning conductance-based models: construction and analysis of databases of model neurons. Journal of neurophysiology, 90(6), 3998–4015. doi:10.1152/jn.00641.2003

Pushpalatha, Z. V., & Konadath, S. (2016). Auditory brainstem responses for click and CE-chirp stimuli in individuals with and without occupational noise exposure. Noise & health, 18(84), 260. doi:10.4103/1463-1741.192477

Ratté, S., Lankarany, M., Rho, Y.-A., Patterson, A., & Prescott, S. A. (2015). Subthreshold membrane currents confer distinct tuning properties that enable neurons to encode the integral or derivative of their input. Frontiers in cellular neuroscience, 8, 452 %@ 1662-5102. doi:10.3389/fncel.2014.00452

Rossant, C., Goodman, D. F., Fontaine, B., Platkiewicz, J., Magnusson, A. K., & Brette, R. (2011). Fitting neuron models to spike trains. Frontiers in neuroscience, 5, 9. doi:10.3389/fnins.2011.00009

Rumbell, T. H., Draguljić, D., Yadav, A., Hof, P. R., Luebke, J. I., & Weaver, C. M. (2016). Automated evolutionary optimization of ion channel conductances and kinetics in models of young and aged rhesus monkey pyramidal neurons. Journal of computational neuroscience, 41(1), 65–90 %@ 0929-5313. doi:10.1007/s10827-016-0605-9

Rumelhart, D. E., Hinton, G. E., & Williams, R. J. (1985). Learning internal representations by error propagation. Retrieved from

Santana, R., Bielza, C., & Larrañaga, P. (2011). Optimizing brain networks topologies using multi-objective evolutionary computation. Neuroinformatics, 9(1), 3–19. doi:10.1007/s12021-010-9085-7

Schneider, S., Igel, C., Klaes, C., Dinse, H. R., & Wiemer, J. C. (2004). Evolutionary adaptation of nonlinear dynamical systems in computational neuroscience. Genetic Programming and Evolvable Machines, 5(2), 215–227. doi:10.1023/B:GENP.0000023689.70987.6a

Schrimpf, M., Kubilius, J., Hong, H., Majaj, N. J., Rajalingham, R., Issa, E. B., … Schmidt, K. (2018). Brain-score: Which artificial neural network for object recognition is most brain-like? bioRxiv, 407007. doi:10.1101/624239

Schrimpf, M., Kubilius, J., Lee, M. J., Murty, N. A. R., Ajemian, R., & DiCarlo, J. J. (2020). Integrative Benchmarking to Advance Neurally Mechanistic Models of Human Intelligence. Neuron. doi:10.1016/j.neuron.2020.07.040

Schröder, C., Klindt, D., Strauss, S., Franke, K., Bethge, M., Euler, T., & Berens, P. (2020). System Identification with Biophysical Constraints: A Circuit Model of the Inner Retina. bioRxiv.

Speiser, A., Yan, J., Archer, E. W., Buesing, L., Turaga, S. C., & Macke, J. H. (2017). Fast amortized inference of neural activity from calcium imaging data with variational autoencoders. Paper presented at the Advances in Neural Information Processing Systems.

Sutskever, I., Vinyals, O., & Le, Q. V. (2014). Sequence to sequence learning with neural networks. Paper presented at the Advances in neural information processing systems.

Svensson, C.-M., Coombes, S., & Peirce, J. W. (2012). Using evolutionary algorithms for fitting high-dimensional models to neuronal data. Neuroinformatics, 10(2), 199–218. doi:10.1007/s12021-012-9140-7

Szatko, K. P., Korympidou, M. M., Ran, Y., Berens, P., Dalkara, D., Schubert, T., … Franke, K. (2020). Neural circuits in the mouse retina support color vision in the upper visual field. Nature communications, 11(1), 1–14 %@ 2041-1723. doi:10.5281/zenodo.3760607

Teeter, C., Iyer, R., Menon, V., Gouwens, N., Feng, D., Berg, J., … Hawrylycz, M. (2018). Generalized leaky integrate-and-fire models classify multiple neuron types. Nature communications, 9(1), 1–15 %@ 2041-1723. doi:10.1038/s41467-017-02717-4

Tsodyks, M. V., & Markram, H. (1997). The neural code between neocortical pyramidal neurons depends on neurotransmitter release probability. Proceedings of the National Academy of Sciences, 94(2), 719–723. doi:10.1073/pnas.94.2.719

Wang, X.-J., Hu, H., Huang, C., Kennedy, H., Li, C. T., Logothetis, N., … Tsao, D. (2020). Computational neuroscience: A frontier of the 21st century. National Science Review, 7(9), 1418–1422. doi:10.1093/nsr/nwaa129

Weaver, C. M., & Wearne, S. L. (2006). The role of action potential shape and parameter constraints in optimization of compartment models. Neurocomputing, 69(10-12), 1053–1057. doi:10.1016/j.neucom.2005.12.044

Webb, S., Golinski, A., Zinkov, R., Siddharth, N., Rainforth, T., Teh, Y. W., & Wood, F. (2018). Faithful inversion of generative models for effective amortized inference.

Wright, N. C., & Wessel, R. (2017). Network activity influences the subthreshold and spiking visual responses of pyramidal neurons in the three-layer turtle cortex. Journal of neurophysiology, 118(4), 2142–2155 %@ 0022-3077. doi:10.1152/jn.00340.2017

Yin, W., Schütze, H., Xiang, B., & Zhou, B. (2016). Abcnn: Attention-based convolutional neural network for modeling sentence pairs. Transactions of the Association for Computational Linguistics, 4, 259–272 %@ 2307-2387X. doi:10.1162/tacl_a_00097

